# Deleterious Mutation Accumulation and the Long-Term Fate of Chromosomal Inversions

**DOI:** 10.1101/606012

**Authors:** Emma L. Berdan, Alexandre Blanckaert, Roger K. Butlin, Claudia Bank

## Abstract

Chromosomal inversions contribute widely to adaptation and speciation, yet they present a unique evolutionary puzzle as both their allelic content and frequency evolve in a feedback loop. In this simulation study, we quantified the role of the allelic content in determining the long-term fate of the inversion. Recessive deleterious mutations accumulated on both arrangements with most of them being private to a given arrangement. This led to increasing overdominance, allowing for the maintenance of the inversion polymorphism and generating strong non-adaptive divergence between arrangements. The accumulation of mutations was mitigated by gene conversion but nevertheless led to the fitness decline of at least one homokaryotype under all considered conditions. Surprisingly, this fitness degradation could be permanently halted by the branching of an arrangement into multiple highly divergent haplotypes. Our results highlight the dynamic features of inversions by showing how the non-adaptive evolution of allelic content can play a major role in the fate of the inversion.

**Author Summary:** A chromosomal inversion is a segment of the chromosome that is flipped (inverted arrangement) relative to the normal orientation (standard arrangement). Such structural mutations may facilitate evolutionary processes such as adaptation and speciation, because reduced recombination in inverted regions allows beneficial combinations of alleles to behave as a “single unit”. This locally reduced recombination can have major consequences for the evolution of the allelic content inside the inversion. We used simulations to investigate some of these consequences. Inverted regions tended to accumulate more deleterious recessive mutations than the rest of the genome, which decreased the fitness of homokarotypes (individuals with two copies of the same arrangement). This led to a strong selective advantage for heterokaryotypes (individuals with one copy of each arrangement), maintaining the inversion polymorphism in the population. The accumulation of deleterious mutations also resulted in strong divergence between arrangements. We occasionally observed an arrangement that diverged into a small number of highly differentiated haplotypes, stopping the fitness decrease in homokaryotypes. Our results highlight the dynamic features of inversions by showing how the evolution of allelic content can greatly affect the fate of an inversion.

## Introduction

Chromosomal inversions are large-scale structural mutations that may encompass millions of nucleotides and cause them to segregate together as a single unit due to repressed recombination. A surge of interest in inversions over the last 20 years has shown that inversions occur in a wide variety of taxa [1-3], are often found to have facilitated evolutionary processes such as adaptation and speciation [3-7], and are frequently under balancing selection [7]. However, we lack a solid understanding of how inversions themselves evolve and which factors determine their fate. Critically, inversions are dynamic and behave in qualitatively different ways from single-nucleotide polymorphisms (SNPs), since both their allelic content and their frequency can change over time. Incorporating this concept better into evolutionary theory will improve our ability to explain and predict the evolution of inversions in natural populations [8-11].

A key feature of inversions, and large structural variants in general, is that selection acts at multiple levels. There is direct selection on the inversion itself as the breakpoints alter the DNA sequence. The allelic content of the arrangements is also under selection, which generates indirect selection at the level of the inversion through linkage disequilibrium. As a consequence of this indirect component, selection on inversions may be overdominant due to the presence of recessive deleterious alleles, unique to each arrangement [12].

Another key feature governing the evolution of inversions is the reduction in effective recombination between the standard (S) and inverted (I) arrangements. Recombination proceeds normally in both homokaryotypes (II and SS). However, in heterokayotypes (IS), single crossovers can lead to unbalanced chromosomes and therefore inviable gametes (but see [13] for other mechanisms of recombination repression). Thus, only gene conversion and double crossovers (in larger inversions) contribute to gene flux (i.e. genetic exchange between arrangements [14]), although recent studies have demonstrated that gene conversion occurs at normal or higher rates in inverted regions [15, 16]. Due to the partial repression of recombination, the arrangements behave like independent populations that exchange migrants. Thus, the arrangements suffer from a reduced population size when compared to the rest of the genome; within each arrangement, selection is less effective and genetic drift stronger. This effect is expected to be weak when an arrangement is at intermediate or high frequency but strong when it is rare [10, 17].

This pseudo-population-substructure only affects the inverted region and affects both standing genetic variation and the fate of new mutations. In particular, the decrease in effective population size mentioned above leads to a reduction in the efficacy of purifying selection, making the two arrangements more vulnerable to the maintenance and possible fixation of deleterious mutations. This expected overabundance of deleterious alleles has been reported in the literature across several taxa such as seaweed flies *Coelopa frigida* [18], *Drosophila melanogaster* [8, 19-22], and *Heliconius* butterflies [23].

In the theoretical literature, the role of recessive deleterious mutations has been addressed previously, mainly regarding the invasion of an inverted arrangement [24-26]. However, the long-term consequences of the reduction in efficacy of purifying selection have not been explored. This is of importance because the efficacy of selection is governed by the frequencies of the different karyotypes (II, IS, and SS). In turn, the allelic content of the inverted and standard arrangements determines their marginal fitness and therefore the frequencies of the different karyotypes. This creates a dynamic feedback loop between the frequency and the allelic content of the arrangements, which has to date received little attention in the literature. The effect is not included, for example, in the influential coalescent models of Navarro et al. [10] and Guerrero et al. [17] where arrangement frequencies are determined solely by direct selection on the inversion or indirect selection due to inclusion of locally-adapted alleles [as in 27].

Here we explore the effects of this feedback loop by modelling how the allelic content of an inversion evolves during its lifetime and significantly impacts its long-term fate. Using Slim v2.6 [28], a forward simulation program, we quantify changes in the allelic content of the inverted region over time and elucidate the role of gene conversion in preventing the accumulation of recessive deleterious mutations. We find that the minority arrangement, which experiences the stronger decrease in population size, accumulates mutations rapidly, leading to a swift decline in the fitness of the corresponding homokaryotype. In smaller populations, this process also occurs in the majority arrangement, potentially resulting in a balanced lethal system. We identify a mechanism that can stop the fitness degradation of homokaryotypes, which we term ‘haplotype structuring’. We discuss how our theoretical predictions can be validated empirically, and highlight the relevance of our results to other scenarios of low recombination.

## Results

### Simulations

We modeled an isolated population of diploid individuals at initial mutation-selection balance using SLiM v2.6 [28]. We simulated a population of N=25,000 (with a subset of simulations run for N=5,000) diploid individuals. The genome consisted of three chromosomes of 1Mb, 300 kb of which were coding regions where allelic content was simulated. The allelic content of the rest of the chromosome was not simulated to alleviate the computational load, although recombination could occur anywhere. Coding regions were modelled as 50 kb segments, separated from each other by 100 kb of non-coding regions (i.e. areas where allelic content was not simulated).

To calibrate our model, we chose parameter estimates inspired by *Drosophila melanogaster* [29-31]. In our model, mutations happened at a rate of μ=8.4 × 10^−9^ per bp per generation [32]. All simulated mutations were deleterious (s < 0), recessive, only occurred in coding regions, and affected individual fitness multiplicatively. The magnitudes of fitness effects of deleterious mutations (|s|) were drawn from a Gamma distribution Γ (α=0.5, β=100). To reduce computation time we did not simulate neutral mutations but 5% of *de novo* mutations were effectively neutral (i.e. |s| <1/(2N). Overall recombination rate was defined as the sum of the rate of single crossovers (CO, ρ = 3.0 × 10^−8^ per base pair per meiosis [29, 30]) and gene conversion (GC, γ =1.8 × 10^−8^ per base pair per meiosis [31] for the rate of initiation of a gene conversion event) and corresponded to the rate of initialization of a recombination event. This overall rate was constant along the genome and for all karyotypes. However, the success of recombination initialization differed between genomic regions and karyotypes. We use the term effective recombination rate to describe the difference in realized events between karyotypes due to crossover suppression in the inverted region in heterokaryotypes. It should be noted that SLiM (in its 2.6 version) did not allow for the possibility of double crossover events. Gene conversion track length followed a Poisson distribution with parameter λ = 500 bp [31]. As recombination is generally restricted to females in *D. melanogaster* but occurs in all individuals in our simulation, we divided the overall recombination rate by 2 (and therefore r = (ρ + γ)/2), resulting in r = 2.4 × 10^−8^ per base pair per meiosis.

Simulation with these parameters was not feasible because of the extremely large computational burden. To reduce computation time while maintaining the same evolutionary scenario, we used the common practice of rescaling parameters so that evolutionary processes happened at an accelerated rate (see for example [33]). A recent paper showed that such rescaling may fail to represent the original population genetics accurately when the product of 2Ns is very large [34]. However, this should not be an issue in our simulations as we remain in the parameter space where using rescaled parameters should not significantly affect the genetic diversity of the population. We thus downscaled both population size and genome length by a factor 10 and upscaled the remaining parameters so that 2NμL, 2Ns, 2NrL, λ/L (with L the length of the genome) remained constant.

Following a burn-in of 500,000 generations to ensure that mutation-selection-drift equilibrium was attained, we assumed that an inversion occurs in a random haplotype (i.e. the random haplotype becomes the inverted arrangement and the remaining haplotypes become the standard arrangement). The inversion occurred between two given loci on chromosome one and encompassed 30% of the chromosome and 10% of the genome. In order to ensure that a reasonable proportion of new inversions remained polymorphic for long enough to observe the effects of deleterious mutations, we assumed that the inversion provided a small heterozygote advantage s_HET_ =0.003 or 2Ns_HET_=150. We followed the fate of the newly introduced inverted arrangement over the next 500,000 generations or until the loss of the inversion polymorphism. We recorded the fitness distribution of the various karyotypes and the inversion frequency over time. For a given haplotype, 100 replicates were used to estimate the invasion probability, both with and without gene conversion. We performed the same analysis for 200 haplotypes from 100 random individuals. In addition to the 200 randomly chosen haplotypes, we also considered the fate of the four fittest and four least fit haplotypes (see Figure S1 for how this choice affected the mutational load of the inversion haplotype).

To further explore the parameter space, we performed additional simulations for the four fittest haplotypes. In order to ascertain the effect of s_HET_ on the fate of the inversion we investigated a range of other heterozygote advantages: s_HET_ = 0, s_HET_ = 0.0003 or 2Ns_HET_ = 15, and s_HET_ = 0.006 or 2Ns_HET_ = 300. To explore the effect of GC, we included 9 additional initiation rates of GC (equally distributed between 0 and 1.8 × 10^−8^ per base pair per meiosis). We also considered an inversion encompassing 20% of the genome, to explore the role of the size of an inversion on its fate. Finally, we also considered a smaller population size (N=5,000). All SLiM scripts, analysis scripts, and the seeds used to run the simulations are available at https://gitlab.com/evoldyn/inversion/wikis/home.

### The Fate of the Inversion

We first quantified the fate of the inverted arrangement, with and without the presence of gene conversion, over the short-term (i.e., if the polymorphism was maintained over the first 10,000 generations *versus* fixation or loss) and long-term (i.e., if the polymorphism was maintained over >500,000 generations *versus* fixation or loss). Gene conversion had little to no effect on the short-term fate (Figure 1a) of the inverted arrangement but increased the probability that the inversion was fixed or lost in the long term (Figure 1b). Without GC, the long-term fate of the inversion was decided within the initial ~60,000 generations after appearance of the inversion (Figure 1f; no losses were observed after generation 58,620). At high GC rates, this was no longer true: even if the inverted arrangement successfully invaded, a risk of losing the polymorphism through genetic drift remained (Figure 1d). This occurs when the GC rate is high enough to partly compensate for the lack of crossing over in heterokaryotypes, which partially erases the pseudo-population substructure created by the inversion. At high rates of GC, the mutational load of the majority arrangement, usually the standard, remains low through two processes. First, purifying selection remains effective in the majority arrangement due to its high frequency. Second, mutations spread between arrangements and thus neither contribute to fitness differences between the karyotypes nor impact the fate of the inversion. Under soft selection, i.e., when there are always enough offspring produced to reach carrying capacity, fitness is relative. Therefore, the fixation of deleterious mutations in the whole population does not count towards the mutational load. The high marginal fitness of the majority arrangement, due to this effective removal of deleterious alleles, increases its frequency making fixation through genetic drift more likely, which results in the loss of the inversion polymorphism.

**Figure 1.**
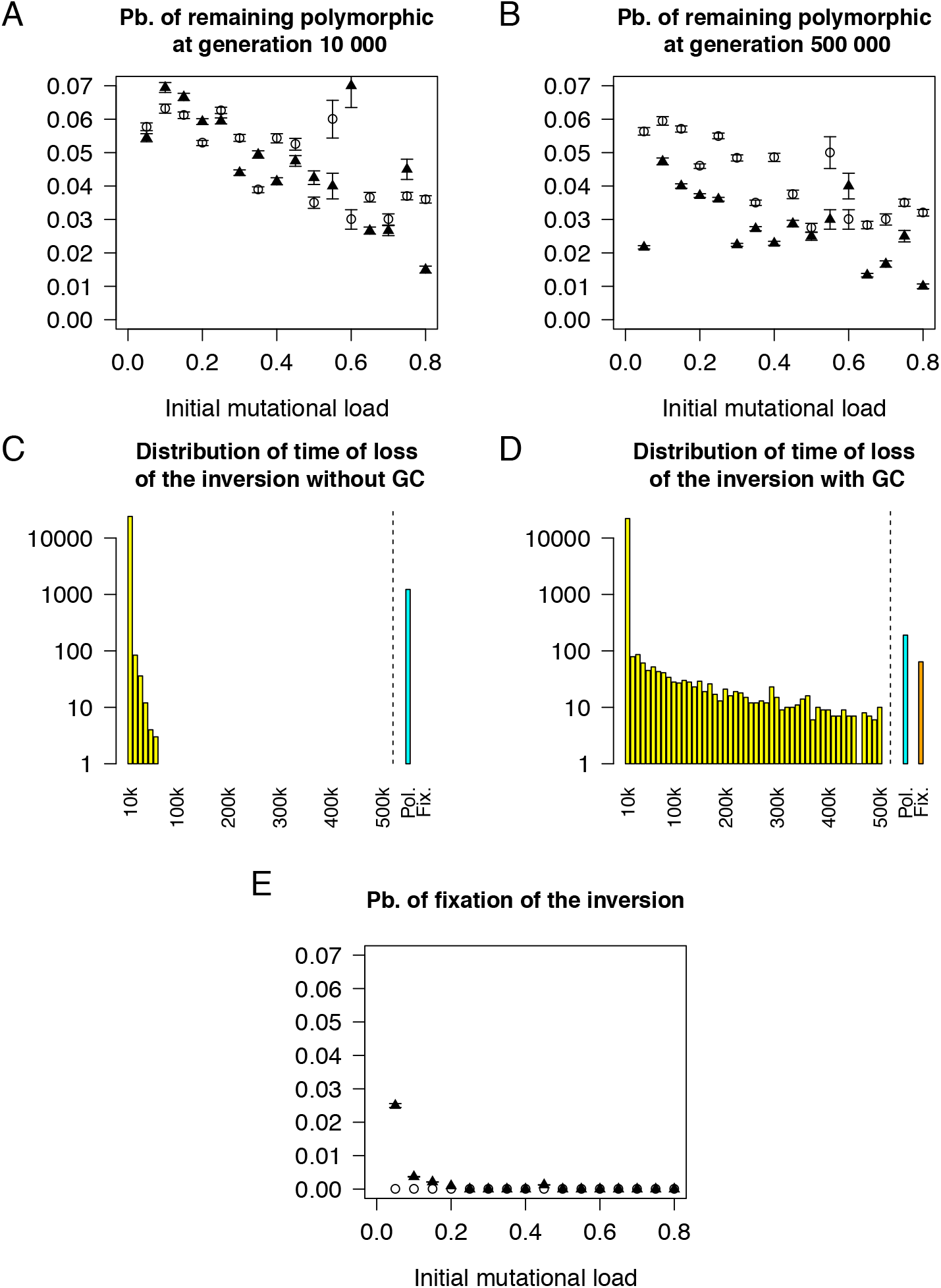
Gene conversion increases the probability that an inversion is fixed or lost. (A) Probability of the inversion being polymorphic at generation 10,000 as a function of the mutational load in the presence (filled) and absence of GC (empty). (B) Probability of the inversion remaining polymorphic at generation 500,000 as a function of the mutational load in the presence (filled) and absence of GC (empty). (C) Distribution of the time of loss of the inversion in the presence of GC. Simulations where the inversion remained polymorphic (cyan) or fixed (orange) are indicated specifically. (D) Distribution of the time of loss of the inversion in the absence of GC. Simulations where the inversion remained polymorphic (cyan) or fixed (orange) are indicated specifically. (E) Probability of fixation of the inversion as a function of the mutational load in the presence (filled) and absence of GC (empty).

Nei and colleagues postulated that an inverted arrangement should be able to spread in a population without additional selective advantage only if it captures a haplotype with low mutational load compared to the rest of the population [24]. This is because inversions originate in a single haplotype; therefore, any inversion homokaryotype (II) will be homozygous for all deleterious recessive mutations present in the original haplotype. Standard homokaryotypes (SS) do not suffer from their mutational load because on average they are homozygous for very few deleterious recessive mutations. Thus, only a few inversion homokaryotypes (II) have a fitness equal to or higher than the mean fitness of the standard homokaryotypes (SS) (Figure S2). In agreement with Nei’s analytical results, we also recovered this pattern in the presence of *de novo* mutation (Nei only considered existing standing genetic variation): we observed fixation of the inverted arrangement when the inversion occurred in a haplotype with a low mutational load (Figure 1e). In the absence of any initial heterozygote advantage (s_HET_=0), both invasion (with probability 0.0082) and fixation (with probability 0.003) were possible, although extremely rare. In addition, we were able to determine that the presence of gene conversion, a lower s_HET_ value (Figure S3), and a smaller population size (N=5,000 Figure S4) all increased the probability of fixation of the inverted arrangement, given invasion has been successful). This is because fixation is only possible if the fitness of the inverted homokaryotype remains similar to the fitness of the heterokaryotype, requiring a low mutational load of the inverted arrangement. In line with this, if heterokaryotype advantage (caused by deleterious or beneficial mutations), i.e. balancing selection, is the driving evolutionary force, fixation will not occur.

### Mutation Accumulation Occurs Inside Chromosomal Inversions

Our results reveal that the content of both the inverted and standard arrangements can change dramatically through the accumulation of recessive deleterious mutations (Figure 2). Generally, the fitness dropped more steeply in the inverted arrangement, but this pattern was reversed when the inversion occurred in a high-fitness haplotype and the inverted arrangement became the majority arrangement. Importantly, whenever the inversion invaded, both arrangements suffered a decrease in both effective population size and effective recombination rate. This decrease in effective recombination rate is due (1) to the absence of crossing over between arrangements and (2) to the reduction in effective population size for each arrangement, leading to a reduction in effective recombination rate within homokaryotypes. This had two important consequences. First, most new mutations remained private to the arrangement they occurred in. Second, recessive deleterious mutations accumulated in the arrangements (Figure 2b,d,f). This accumulation process was unaffected by the strength of the added heterokaryotype advantage (or its presence) in our model (Figure S3). The size of the inversion did not change this process qualitatively although the larger inversion accumulated slightly more mutations (per kb) in the major arrangement in the absence of GC (Figure S5). Accordingly, each arrangement experienced a process similar to Muller’s ratchet, which is the step-wise stochastic loss of haplotypes with the lowest mutational load in the absence of sufficient recombination [35-40]. Despite the accumulation of deleterious mutations, the inversion remained in the population due to the increasing heterokaryotype advantage. This is sometimes referred to as associative overdominance which is caused by linkage disequilibrium between the inversion and alleles within it that confer heterozygote advantage. Both overdominant as well as recessive deleterious alleles may contribute to this phenomenon [3, 9, 25]. In our model, associative overdominance is generated by the presence of private recessive deleterious alleles at different loci in the two arrangements. The inversion polymorphism is therefore maintained by genic selection where inversions act as neutral vehicles of selected alleles, *sensu* Wasserman [26, 41]. Thus, deleterious mutation accumulation provides the raw material upon which genic selection acts, leading to the maintenance of the inversion polymorphism.

**Figure 2.**
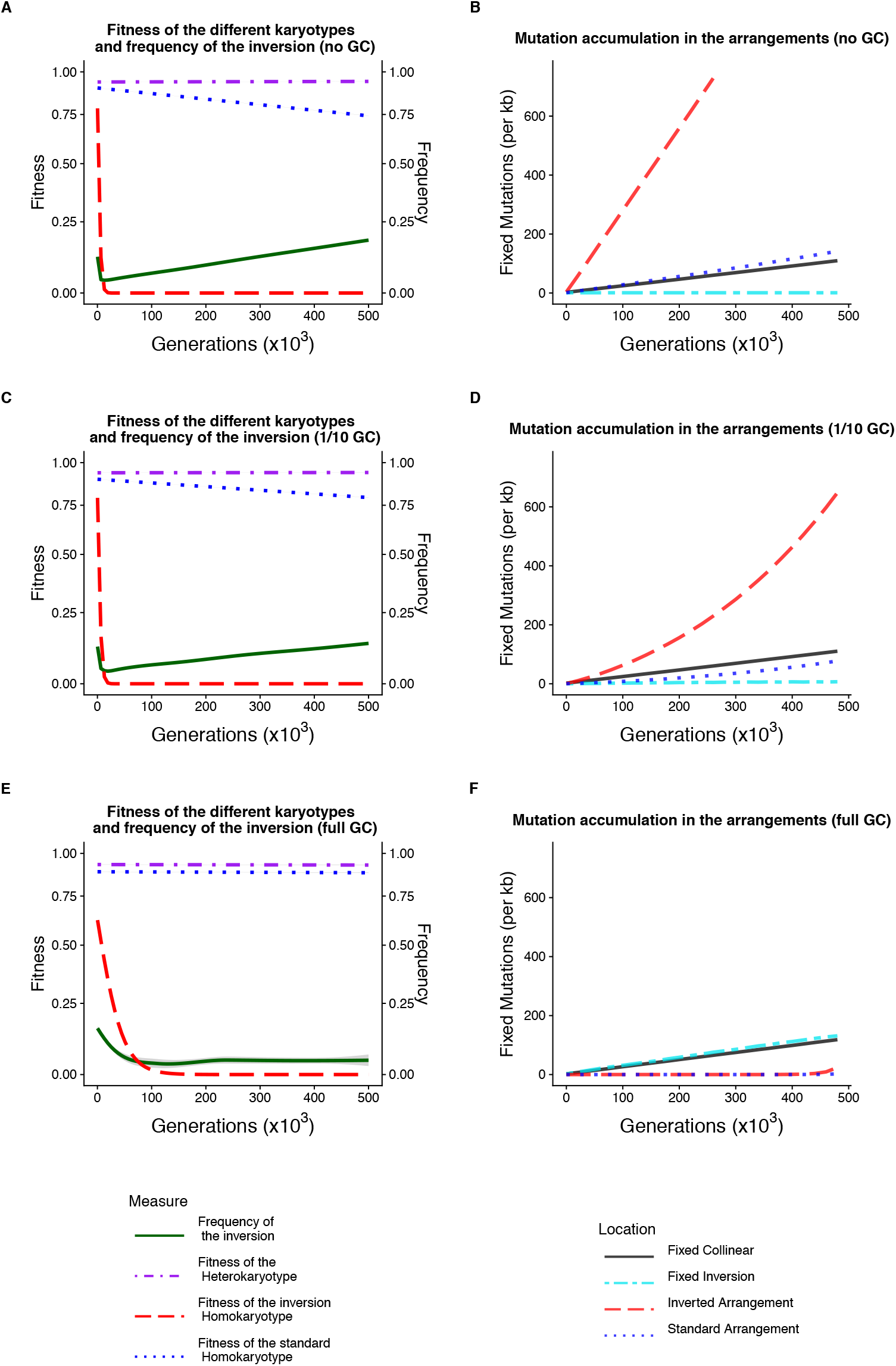
Fitness decay of the homokaryotypes and accumulation of mutations in the different arrangements (A,C,E). Fitness of the different karyotypes for the inversion and frequency (green) of the inversion over 500,000 generations (starting at generation 200 after introduction) following the introduction of the inversion under (A) a scenario with no gene conversion, (C) a scenario with 1/10 of the *D. melanogaster* gene conversion rate, and (E) a scenario with the *D. melanogaster* gene conversion rate. (B,D,F) Corresponding cumulative distribution of fixed mutations per kb in the inverted arrangement (red), the standard arrangement (blue), the inverted region (turquoise), and in the collinear region (black) depending on the generation when the mutation appears. Results were obtained from 1,000 replicates where we only display successful maintenance of the inversion polymorphism (5 cases with a high rate of GC, 60 cases with 1/10 of the previously used GC rate GC, and 61 cases without GC).

The level of gene flux (i.e. genetic exchange between the two arrangements), determined solely by gene conversion in our model, is a key factor in determining the allelic content of the arrangements. As illustrated in Figure S6, both the number of mutations and the mutational load of a given arrangement decrease in an exponential-like fashion with an increase in GC rate. However, the major and minor arrangements were differentially affected. While the minority arrangement always accumulated mutations at a much faster rate than the majority arrangement, the addition of gene conversion to the model decreased the number of deleterious mutations in both arrangements (Figure 2b,f). On average, both the majority and minority arrangement had > 20 times more mutations in the absence of GC (majority arrangement: 23x, 95% confidence interval (CI) from bootstrapping: 18.3-29.0; minority arrangement: 28x, 95% CI 15.3-53.4). Yet the fitness of the two arrangements was not equally affected by high GC rates. The fitness of the majority homokaryotype was scarcely affected by mutation accumulation (because a small decrease in its population size resulted in a slightly larger mutational load), whereas the fitness of the minority homokaryotype decreased to ~0 (<10^−3^). Non-zero GC rates allowed both mutations and ancestral alleles to move between arrangements and fix in the whole population, which reduced divergence between arrangements (see below) and aided the purging of deleterious mutations. We only observed a single instance whereby purging of deleterious mutations allowed the fitness of an arrangement to recover successfully from close to 0 (see Supplemental Text). At low GC rates, the global fixation rate of mutations within the inverted region (i.e. mutations that spread across arrangements) was reduced (see cyan line, Figure 2b,d). However, at sufficiently high GC rates, mutations could spread across arrangements and fix in the whole population at a similar rate to the collinear genomic regions (see cyan line, Figure 2f). Thus, the mutational load of the individual arrangements remains lower at high GC rates, but ancestral alleles can be irreversibly lost from the whole population.

The population size also has a strong impact on the long-term fate of the inversion. In larger populations, mutation accumulation was either stopped or bypassed (see Section *Appearance of Haplotype Structuring* below) and only the minority homokaryotype became inviable (defined here as having an average relative fitness < 0.001). This was always the case at high GC rates and almost always in its absence (1218/1227 99.3% of completed runs). In small populations, weaker purifying selection led to an additional evolutionary outcome where both homokaryotypes became inviable. In this case, only heterokaryotypes contributed to subsequent generations. This long-term outcome was observed both in the absence of GC (56/56 test cases in which the inversion polymorphism remained) and at high rates of gene conversion (10/15 test cases in which the inversion polymorphism remained). Thus, at small population sizes, an inversion polymorphism may trigger the development of a balanced lethal system, various cases of which have been observed in nature [42-47].

### Mutation accumulation causes strong divergence between arrangements

Whenever the inverted arrangement invaded, mutation accumulation within each arrangement resulted in fixed differences between the inverted and standard arrangement (Figure 3a,b). Unsurprisingly, more fixed differences accumulated in the absence of gene conversion (average number of fixed mutations without GC: 4,609 ± 7) than in its presence (average number of fixed mutations with GC: 182 ± 2). This strong between-arrangement divergence was reflected in high overall F_ST_ values between arrangements within the inverted region, compared with little divergence across the rest of the chromosome (Figure 3). Notably, no beneficial mutations are necessary for the buildup of the between-arrangement divergence. To better understand the role of purifying selection, we can separate the deleterious mutations into two categories: effectively neutral mutations (i.e. |s| <1/(2N)) and deleterious mutations. In our simulations, about 5% of new deleterious mutations are effectively neutral considering the total population size. If purifying selection is a potent force, we expect most fixed mutations to be effectively neutral. We find that purifying selection in large populations was relatively effective in collinear regions as ~50% of the fixed mutations were effectively neutral (Figure S7). However, within the two arrangements, the effectiveness of purifying selection was strongly decreased, particularly in the minor arrangement. This is evidenced by the proportion of effectively neutral fixed mutations in simulations without GC (majority arrangement: 46.1% ± 0.1%; minority arrangement: 5.2% ± 0.03%). The presence of GC altered the number of fixed mutations within arrangements (see above) but barely affected the proportion of effectively neutral fixed mutations (majority arrangement: 43.6% ± 0.9%; minority arrangement: 5.4% ± 0.1%). Surprisingly, some fixed mutations were very strongly deleterious (Figure S8). Both the strong within-arrangement divergence and the observation of less effective purifying selection support the interpretation of an inversion as a region of the genome that experiences population-substructure.

**Figure 3.**
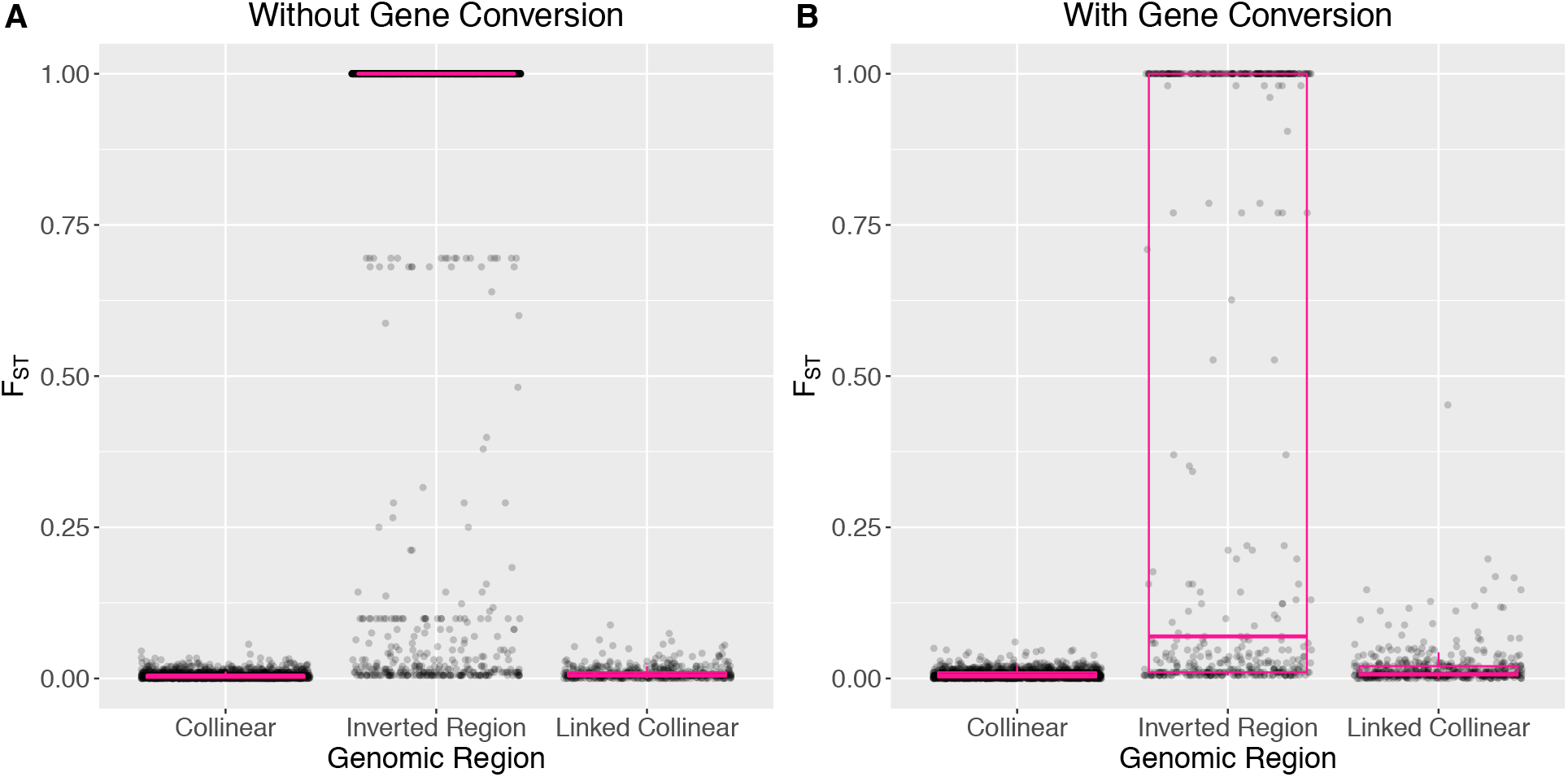
Divergence between karyotypes in the collinear, inverted, and linked regions. Linked regions are on the same chromosome as the inverted region but not within it. Each dot represents a single SNP and boxplots are overlain in pink. (A). F_ST_ without gene conversion, (B). F_ST_ with gene conversion.

### Appearance of haplotype structuring

The fitness degradation of one or both arrangements that we describe above was occasionally halted by a mechanism we term *haplotype structuring* if GC rate was low enough (Figure S6). When haplotype structuring occurred, the subpopulation of one arrangement split into two or more divergent haplotype clusters that carried partially complementary sets of deleterious recessive alleles (see Figure 4 & 5). Here, homokaryotypes with two divergent haplotypes that each have a high mutational load are still relatively fit (e.g. I_j_I_k_ and S_j_S_k_) because deleterious mutations are masked when divergent haplotypes are paired. Notably, this is equivalent to what is happening in heterokaryotypes (IS). Homokaryotypes with similar haplotypes (e.g. I_j_I_j_ or S_j_S_j_) tend to be inviable because the mutational load is no longer masked. This means that the fitness distribution of a given homokaryotype (e.g. II) has two modes; one corresponding to extremely unfit individuals and the other to relatively fit ones (see Figure 5 for a schematic). Thus, a signature of haplotype structuring in a given arrangement is that the fitness of the corresponding homokaryotypes shifts from a unimodal to a bimodal distribution (Figure S9). We also recover this result in the absence of direct heterozygote advantage for the inversion (s_HET_ =0). Figure S10 depicts an outcome similar to Figure 4B: haplotype structuring in the major arrangement. When haplotype structuring occurs, the expected equilibrium frequency of the inversion tends to be close to 0.5, due to the large fitness advantage of the heterokaryotypes over the homokaryotypes. However, the expected equilibrium frequency still depends on the marginal fitness of the two homokaryotypes (Figure 4B, 4D), and will only be equal to 0.5 if the mutational load is, and remains the same in both arrangements (the balanced lethal case is one such example, Figure 4A).

**Figure 4.**
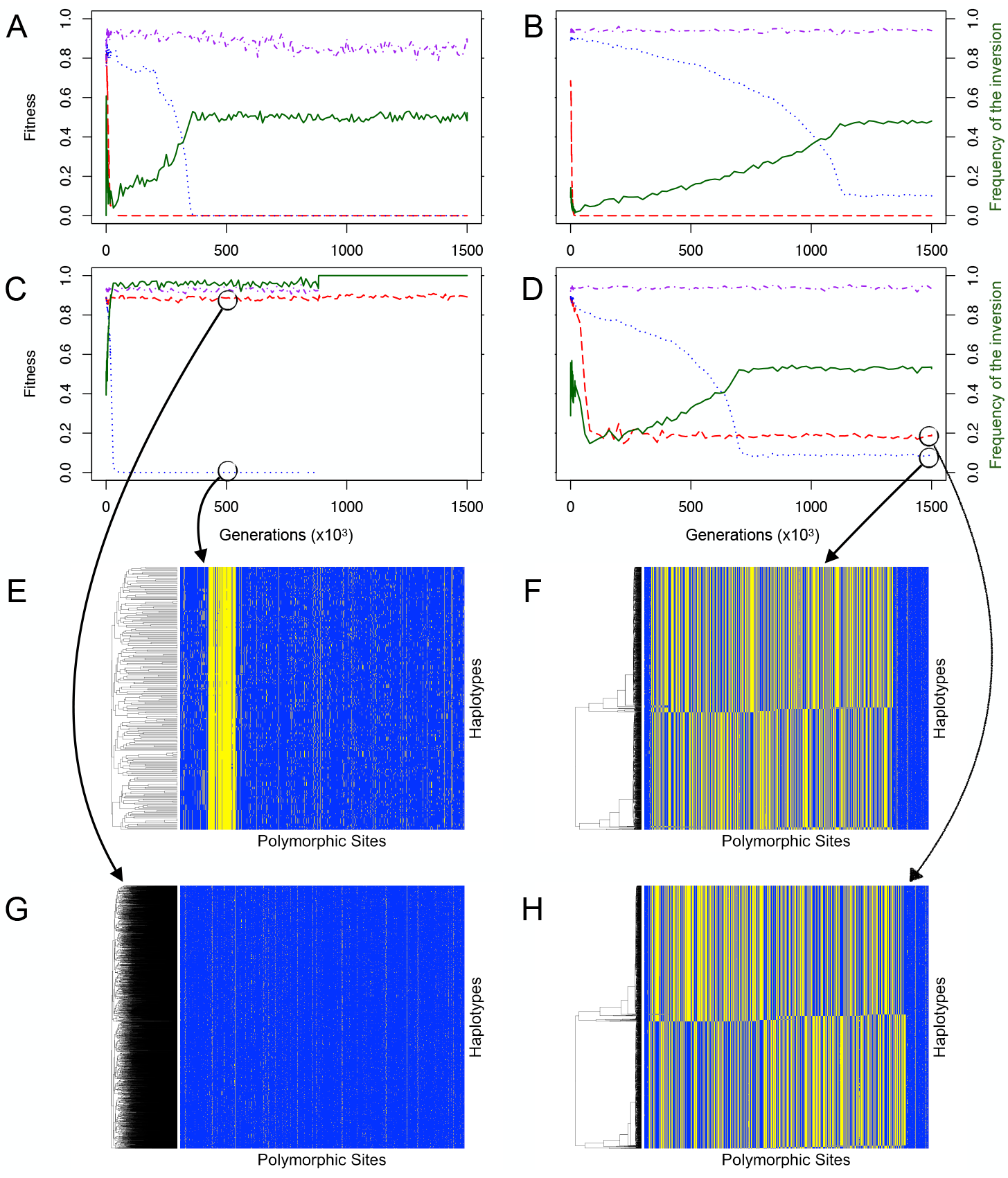
Different evolutionary outcomes (A-D) and allelic content of the arrangements (E-H). (A-D) represent the fitness of the different karyotypes as well as the frequency of the inversion for all 4 outcomes. Fitness of the standard homokaryotype is given by the dotted blue line, of the inverted homokaryotype by the red dashed line and of the heterokaryotype by the dash-dotted purple line. The frequency of the inversion is given by the solid green line. A) Balanced lethals, B) inverted homokaryotypic is inviable, standard homokaryotype remains viable through haplotype structuring: C) inverted homokaryotype is viable, standard homokaryotype is inviable until the inversion fixes, D) haplotype structuring in both the inverted and standard arrangements. (E-H) Allelic content of the inversion, each horizontal line represents a haplotype in the population and each vertical line represents a genomic locus. Yellow denotes that an individual possesses the derived allele and blue the ancestral one. The black circle indicates where the haplotypes were taken from. E) Mutation accumulation in the minor arrangement, F) haplotype structuring in the standard arrangement, G) purifying selection in the majority arrangement, H) haplotype structuring in the inverted arrangement.

**Figure 5.**
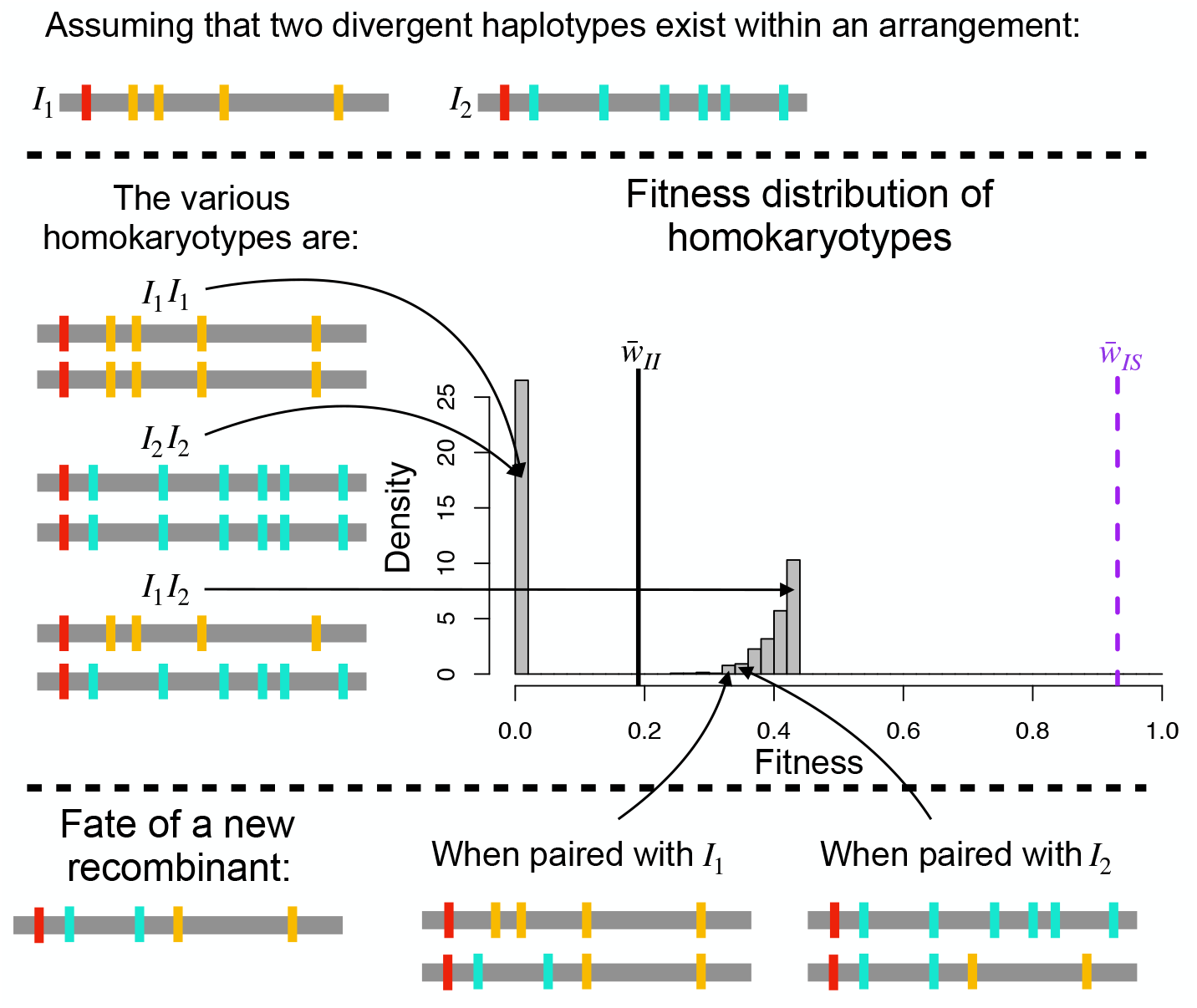
Schematic representation of the consequences of haplotype structuring on the fitness distribution of the homokaryotypes. Red, cyan, and mustard represent deleterious mutations. Homokaryotypic homozygotes have a fitness near 0 while homokaryotypic heterozygotes have a positive fitness, as only the mutations that are fixed in the arrangements (in red) are expressed, while the mutations unique to each haplotype (in mustard and cyan) are masked. This leads to the bimodal distribution of fitness illustrated here. For reference the vertical lines correspond to the mean fitness of heterokaryotypes (dashed purple) and homokaryotypes (black line). Haplotype structuring is stable against recombination as the new recombinant will express both mustard and cyan mutations, leading to a lower fitness, whenever it is associated with either of the two major haplotypes.

Haplotype structuring requires a significant level of within-arrangement diversity. Namely, the mutational load of the segregating haplotypes has to be high to create a large fitness difference between homokaryotype homozygotes (e.g. I_j_I_j_ or S_j_S_j_) and homokaryotype heterozygotes (e.g. I_j_I_k_ or S_j_S_k_), which in turn generates within-arrangement genic selection. Therefore, haplotype structuring is not possible in small populations or at high GC rates. Indeed, we only observed haplotype structuring with GC rates γ ≦ 5.4 × 10^−9^ (Figure S6). At high GC rates, the mutational load of the majority arrangement is not sufficiently large for haplotype structuring to occur and there are not enough copies of the minority arrangement present to create the necessary diversity. Similarly, in small populations, the haplotype diversity necessary for haplotype structuring cannot build up or be maintained because it is overwhelmed by the diversity-reducing force of genetic drift.

The divergent haplotype clusters that result from haplotype structuring are stable and are not disrupted by recombination. This is because recombination between divergent haplotypes creates new haplotypes that expose deleterious recessive mutations to selection when paired with either one of the parental haplotypes. Therefore, any recombinant haplotype is swiftly removed from the population even though its deleterious mutations are not exposed to selection in a heterokaryotype. Haplotype structuring has previously been described by Charlesworth and Charlesworth in a model of a diploid non-recombining population with deleterious recessive mutations [48]. To confirm this similarity, we triggered haplotype structuring in simulations of whole genomes with greatly reduced recombination rates. In these simulations, haplotype structuring was possible across the full range of GC rates we tested as long as crossing-over rates were low (20% or less of our default value, Figure S11). Thus, similar to how heterokaryotype advantage maintains an inversion polymorphism, heterozygote advantage at the level of the haplotype maintains the haplotype polymorphism (i.e. haplotype structuring). Importantly, although haplotype structuring halts the fitness decay of homokaryotypes, mutation accumulation continues.

## Discussion

Chromosomal inversions are dynamic variants that behave in qualitatively different ways from other polymorphisms (SNPs, indels). Specifically, both their allelic content and their frequency change over time, leading to two intertwined levels of evolution. We demonstrate here that the allelic content of an arrangement can degrade via a Muller’s ratchet-like process. While the inversion remains polymorphic in the population, we observe an accumulation of deleterious recessive mutations in one or both of the arrangements, which can result in at least one of the homokaryotypes becoming inviable. In our simulations, this fitness decay is slowed by gene conversion but can only be stopped by haplotype structuring, the appearance of multiple highly-divergent haplotypes within an arrangement. Together, our results imply that inversions observed in nature can be substantially different from the original invader even without the action of directional selection. Furthermore, we predict that they may harbor sub-haplotypes within arrangements that can distort population genetic statistics.

We show that a mutation accumulation process similar to Muller’s ratchet happens within the arrangements that experience a reduced effective recombination rate and a reduced effective population size. These reductions decrease the efficacy of purifying selection resulting in an excess of deleterious mutations within the inverted region compared to the rest of the genome. This relationship between recombination and the efficacy of selection is well documented [49-51]. The increased accumulation of deleterious mutations in polymorphic inversions compared to collinear regions has previously been noted in multiple empirical studies. By crossing within and between populations Butlin and Day showed that a significant proportion of the observed heterokaryotype advantage in seaweed flies (*Coelopa frigida*), could be ascribed to associative overdominance caused by deleterious recessive mutations [18]. A similar result was found in *D. pseudoobscura* where crosses between populations yielded fitter homokaryotypes than crosses within populations [52]. Likewise, in *Drosophila melanogaster,* inversion-carrying chromosomes were more likely to carry lethals than inversion-free chromosomes [20]. Even when excluding lethal mutations homokaryotypes still had significantly lower fitness than heterokaryotypes indicating overdominant mutations [20, 22]. Another study in *D. melanogaster* found that minority arrangements in wild populations contained significantly more p-elements [8]. A follow-up study also found increased numbers of transposable elements (TEs) in low frequency inversions [21]. Here, the authors argued that the rate of back mutation (i.e. removal of TEs) was too high to allow for continued accumulation as predicted under Muller’s ratchet. Other studies have shown that the efficacy of selection is reduced in inversions. In the laboratory, lethal alleles located within inversions in *Drosophila melanogaster* were maintained at similar frequencies for over 100 generations indicating that selection was not effective [19]. Next generation sequencing has allowed more detailed surveys of inversion content. A recent study by Jay *et al.* [23] examined the content of the P supergene in *Heliconius numata* which encompasses two chromosomal inversions. They found an enrichment of non-synonymous relative to synonymous substitutions, negative selection on the arrangements, and a larger proportion of transposable elements compared to the rest of the genome [23]. Overall, these results indicate that mutation accumulation may be a common process in natural inversions, where the types of mutations that are accumulated can vary.

The rate of mutation accumulation differs between the standard and inverted arrangements. The extent of this difference depends on the relative frequency of the two homokaryotypes, as most “genome shuffling” occurs within homokaryotypes. Mutation accumulation is magnified in the minority arrangement as the associated subpopulation experiences a stronger reduction in population size and therefore a lower effective recombination rate (approx. rp^2^, with r - the recombination rate and p - the frequency of the minority arrangement). Moreover, the purging of recessive deleterious mutations is less effective in the minority arrangement as the respective mutations are only exposed to selection in few individuals. Eanes *et al.* developed a model showing that the minority arrangement accumulated more p-elements at lower frequencies and predictions from this model matched empirical data from *D. melanogaster* [8]. Other empirical studies have also illustrated the relationship between arrangement frequency and mutational load [53-55]. Most notably, Tuttle *et al.* examined the 2^m^ allele (an arrangement of an inverted region on chromosome 2) in white-throated sparrow (*Zonotrichia albicollis*), which exists almost exclusively in the heterokaryotypic state [56]. They found that 2^m^ contained an excess of non-synonymous fixed mutations, which is consistent with functional degradation. Here, by revealing the feedback loop between arrangement frequency and mutational load, we present an intuitive reasoning for these observations.

The accumulation of recessive deleterious mutations in the arrangements led to heterokaryotype advantage caused by the masking of recessive mutations. In the theoretical literature, there is a large body of work focusing on the role of recessive deleterious mutations with regard to the invasion of a new inversion [24-26]. This body of work has concentrated on the role of existing standing genetic variation. In contrast, we do not know of theoretical work that has addressed the role of *de novo* deleterious mutations in the long-term maintenance of an inversion polymorphism. In nature, a contribution of deleterious recessive alleles to heterokaryotype advantage has been inferred in seaweed flies [18], but it is unknown whether these mutations predate the inversion itself. Furthermore, similar empirical tests in other taxa remain scarce. As heterokaryotypes are often observed to be fitter than homokaryotypes [57-59], mutation accumulation may commonly play a role in the maintenance of inversion polymorphisms.

In the age of next generation sequencing, the genomic landscape of many inversions is being dissected to elucidate the processes driving inversion evolution [7, 60]. Our work adds to past theoretical results showing that regions of low recombination may accumulate neutral divergence (ex: Navarro et al [10]). Since various natural inversions have been reported to influence adaptive traits, divergence observed between arrangements has often been assumed to be adaptive and/or to predate the inversion itself, whereas the process of deleterious mutation accumulation has received little attention [7, 13]. However, not only adaptation and but also simply drift are able to generate this pattern of diversity in inversions [17]. We partly recover this result: we show, in Figure 3, that it is possible for fixed mutations between different arrangements to be neither adaptive nor predating the inversion. The strong divergence between arrangements that results from deleterious mutation accumulation can produce a similar population genetic signature to that of a cluster of (co-)adapted alleles within an arrangement [61-63].

We were specifically interested in the long-term evolutionary fate of the inversion, when both arrangements were maintained in the population. We identified multiple stable evolutionary outcomes for each arrangement under deleterious recessive mutation accumulation (over 60N generations). They can be divided into three general categories, depending on the mutational load of the arrangement and the fitness of its corresponding homokaryotype.

First, if the mutation accumulation and the associated gradual decrease in homokarypotype fitness continued, then the corresponding homokaryotype eventually became inviable. This often occurred in only the minority arrangement. In this case the polymorphism was maintained but the minority arrangement only appeared in heterokaryotypes. When the corresponding homokaryotypes of both arrangements became inviable, only heterokaryotypes contributed to subsequent generations. Thus, the mutation accumulation process shown here is a credible model for the evolution of a balanced lethal system. Our results show that low population size and reduced gene flux favor the evolution of balanced lethality. Several empirical examples of balanced lethal systems associated with structural variants exist. These include multiple overlapping structural variants in crested newts [46], inversions in *Drosophila tropicalis* [43], and translocations (similar to inversions, effective recombination in the translocated regions is also reduced) in multiple genera of plants such as *Isotoma* [44], *Rhoeo* [45], *Gayophytum* [47] and *Oenothera* [42]. Using a mathematical model inspired by the latter system, de Waal Malejit and Charlesworth proposed that the accumulation of deleterious recessive mutations could create sufficient mutational load for the maintenance of translocation heterozygosity in a selfing population, assuming a large enough mutational target [64]. To provide evidence for the evolution of balanced lethal systems through mutation accumulation in structural variants, inference of the demographic history of these populations will be essential in the future.

The second long-term outcome is the maintenance of a highly fit homokaryotype with low mutational load of the corresponding arrangement. This outcome was only observed in the majority arrangement and at high GC rates. Here the mutation accumulation is truly stopped as opposed to the case of haplotype structuring, where the consequences of mutation accumulation are bypassed. While the majority homokaryotype maintains a stable, high fitness, the fitness of the minority homokaryotypes drops to 0. When this occurs, the minority arrangement remains at very low frequency (s_HET_ /(1+ 2s_HET_) if the fitness advantage of the heterokaryotype over the majority homokaryotype is only due to the imposed initial heterozygote advantage). Thus, this outcome is the least stable as the high frequency of the majority arrangement combined with a small fitness difference between heterokaryotypes and majority homokaryotypes facilitates fixation of the majority arrangement.

The third category of long-term stable outcomes involves haplotype structuring in one or both of the arrangements. Haplotype structuring halts the fitness decay of the corresponding homokaryotype but it does not stop the mutation accumulation process. As illustrated in Figure 5, the existence of two (or more) divergent haplotype clusters within an arrangement implies that most mutations will be masked in homokaryotype heterozygotes (e.g. I_j_I_k_ or S_j_S_k_). Similarly to what happens between arrangements, mutations tend to be private to haplotype clusters. Therefore, a subset of homokaryotypes still contributes to the next generation. The fitness consequences of mutation accumulation are merely bypassed due to the recessivity of the deleterious mutations. It is critical to note that haplotype structuring as described here is a within-population mechanism as both drift and selection are required. Whereas the same outcome (divergent haplotypes) may be obtained in separate populations [65], drift should be sufficient to explain this pattern. Thus, haplotype structuring is not expected to evolve in highly structured populations with little migration between them.

Wasserman showed that if the fitnesses of both homokaryotypes are reduced due to the existence of a recombinational load, a heterokaryotype fitness advantage will appear [12]. The recombinational load in the Wasserman model is caused by the existence of multiple divergent haplotypes containing a balanced combination of epistatically interacting alleles. Here, we show that accumulation of deleterious recessive mutations can generate a similar pattern. However, with dominance, this effect is due to a combination of segregational and recombinational load. Thus, recombinational load can be generated without epistasis. In both models, the key element for reduction in homokaryotype fitness is the existence of interactions at the gene level (either intra- or inter-locus) that lead to the formation of a recombinational or/and segregational load for the homokaryotypes.

Haplotype structuring occurs when a continual input of deleterious mutations results in associative overdominance in regions of low recombination, where it increases genetic diversity by maintaining complementary heterozygous haplotypes. Thus, the occurrence of haplotype structuring is not unique to inversions. It can also occur in diploid low-recombination systems with segregation of chromosomes. We were able to reproduce haplotype structuring using simulations with similar conditions but without assuming an inversion, provided there was a strong decrease in crossing-over rate (Figure S11). Using a theoretical model, Gilbert *et al.* recently showed that haplotype structuring can occur in regions of low recombination under quite general conditions, especially if deleterious selection coefficients are of intermediate strength [66]. Importantly, they demonstrated that the pattern of increased diversity caused by associative overdominance (likely a result of haplotype structuring) is also sustained with incomplete dominance. Moreover, the predicted pattern of increased diversity was observed in human genomic data [66].

Haplotype structuring has been described previously [48], where the authors modeled the accumulation of deleterious recessive mutations in a diploid, non-recombining, random-mating, sexual population and noted that the population could become crystallized into two divergent haplotypes. Although we recovered the crystallization part of the process, we sometimes observed more than two haplotype clusters (Figure S12). In this case, fitness could be multimodal (Figure S12b) depending on the fitnesses of the different homokaryotype heterozygotes. A larger number of divergent haplotypes increases the average fitness of homokaryotypic individuals because homozygotes (e.g.: I_j_I_j_ S_j_S_j_) are inviable and their proportion (given by: 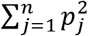, i.e. the sum of all possible homokaryotype homozygotes) decreases as the number of haplotype clusters increases. Therefore, the number of haplotype clusters obtained is the result of a balance between genic selection, which selects for many haplotype clusters, and genetic drift, which reduces the number of haplotype clusters. Once clusters are formed, new recombinant haplotypes are counterselected due to the high number of shared recessive deleterious mutations between a recombinant and a resident haplotype (Figure 5).

Whereas various examples of balanced lethals are known (discussed above), we are not aware of existing empirical evidence for haplotype structuring in inversions. This could be for two reasons. First, compensatory evolution and/or selective sweeps of beneficial mutations within the arrangements could erase haplotype structuring. We are currently not including beneficial mutations in our simulations; adding them to the model would lead to selective sweeps that should reduce the diversity within the (sub)population. Therefore the initial requirement of strongly divergent haplotypes would possibly not be met. Second, the pattern may have remained invisible to date due to the low density of markers available in the past as well as the current common practice of pooled sequencing, which does not reveal haplotypes. Additionally, other aspects of experimental design - for example breeding designs that allow the fitness of offspring of each mating pair to be measured - are necessary to detect the predicted bimodal fitness distribution. Future empirical work could investigate these patterns, testing explicitly for bimodal fitness distributions and for the existence of clusters of haplotypes within arrangements using individual re-sequencing data.

There are several limitations to our study. First, we focus on deleterious mutations. The inclusion of beneficial mutations will affect the invasion process and the probability of the inverted arrangement fixing: the effects of such mutations on an existing polymorphic inversion remain unclear. The spread of a beneficial allele within an arrangement will cause a loss of genetic diversity and the corresponding increase in mutational load could cancel out the initial selective advantage provided by the beneficial mutation. We hope to investigate this in future work. Second, we considered all deleterious mutations to be fully recessive. Incomplete dominance may slow the accumulation of deleterious mutations but is unlikely to stop it. Preliminary work shows that as long as recombination is low enough and selection maintains the structural variant polymorphism, even fully dominant deleterious mutations will accumulate (Gutiérrez-Valencia, pers. comm.). Third, we only consider gene conversion as a mechanism for gene flux between arrangements and not double crossovers. Double crossovers transfer larger tracts of sequence and thus their inclusion will increase gene flux. This would have similar consequences to increasing the GC rate (see Figure S6) and would likely decrease the rate of mutation accumulation and all potential ensuing processes (e.g. haplotype structuring). However, evidence suggests that double crossover rates within inversion heterokaryotypes are reduced compared to rates in homokaryotypes or collinear regions [67-69]. Furthermore, the contribution of double crossovers to gene flux is negligible as long as the size of the inversion is small compared to the inverse of the rate of double strand breaks [70]. Finally, computational limitations prevented us from exploring a wide range of population sizes and inversion sizes. While we do not expect these parameters to alter our qualitative conclusions, it is difficult to predict their quantitative effects.

Our results show that inversions are dynamic variants whose allelic content can evolve and impact their evolutionary fate. We also show that non-adaptive processes in inversions can generate “adaptive-like” signatures. These results stress that the evolution of the allelic content of the inversion should be included in future models and in interpretations of sequence variation in inversions. Our study suggests several particular evolutionary outcomes of inversion evolution, which are potentially also applicable to regions of low recombination. The advent of improved methods for genome assembly should make it possible to determine how often haplotype structuring and balanced lethals occur in nature.

## Materials and Methods

Simulations were implemented in SliM v2.6 [28] (scripts, analysis scripts, and seeds available at https://gitlab.com/evoldyn/inversion/wikis/home)

## Acknowledgements

We thank the Bank lab for support and helpful comments on the study design and the manuscript. We thank the editor and the reviewers for constructive comments on earlier versions of this manuscript. We thank I. Fragata for advice on study design and figures, I. Gordo for valuable advice, and B. Charlesworth, A. Westram and K. Johannesson for helpful comments on the manuscript. E. B. was supported by a Marie Skłodowska-Curie fellowship 704920 – ADAPTIVE INVERSIONS. R.K.B. was supported by the NERC and by ERC Advanced Grant 693030 - BARRIERS. C.B. is grateful for support by EMBO Installation Grant IG4152. A.B. and C.B. were supported by ERC Starting Grant 804569 - FIT2GO.

## Author Contributions

E.B. and R.K.B. conceived of the study. E.B. and A.B. designed the simulations. A.B. wrote the analysis scripts. A.B. and E.B. analyzed the data. C.B supervised the project. All authors interpreted the results and wrote the paper.

## Materials & Correspondence

Requests for material and correspondence can be addressed to E.B and A.B.

## Supplemental Figure Legends

**Figure S1.** Density distribution of the initial mutational load. A) the mutational load in the whole population at the end of the burn-in. B) the mutational load of the inverted arrangement in the haplotypes we selected (200 random plus the 4 best and the 4 worst and one close to the median). C) the mutational load of the inverted arrangement after correcting for the number of simulations done per haplotype. This figure illustrates that we do not always have the same number of simulations for each datapoint in Figure 1.

**Figure S2.** Distribution of the initial relative fitnesses of all 3 karyotypes when an inversion occurs in any haplotype in a population.

**Figure S3.** Effects of the added heterokaryotype advantage. (A) Distribution of the time of loss of the inversion at s_HET_=0. The number of simulations that remained polymorphic (cyan) or fixed (yellow) are indicated specifically to the right of the dashed line. (B) Distribution of the time of loss of the inversion at s_HET_=0.0003. The number of simulations that remained polymorphic (cyan) or fixed (yellow) are indicated specifically to the right of the dashed line. (C) Distribution of the time of loss of the inversion at s_HET_=0.003. The number of simulations that remained polymorphic (cyan) or fixed (yellow) are indicated specifically to the right of the dashed line. (D) Distribution of the time of loss of the inversion at s_HET_=0.006. Simulations that remained polymorphic (cyan) or fixed (yellow) are indicated specifically to the right of the dashed line. (C) Mutation accumulation in the major arrangement under s_HET_=0 (red), s_HET_=0.0003 (green), s_HET_=0.003 (cyan), and s_HET_=0.006 (purple). Each dot represents a single run that ended at generation 500,000. (D) Mutation accumulation in the major arrangement under s_HET_=0 (red), s_HET_=0.0003 (green), s_HET_=0.003 (cyan), and s_HET_=0.006 (purple). Each dot represents a single run that ended at generation 500,000.

**Figure S4.** Distribution of the time of loss of the inversion at different population sizes. For A and B N=25,000 and for C and D N=5,000. A and C show simulations run without gene conversion and C and D show simulations with gene conversion added. All plots show distribution of the time of loss of the inversion. Simulations that remained polymorphic (cyan) or fixed (yellow) are indicated specifically to the right of the dashed line.

**Figure S5.** Mutation accumulation (A,B) and Mutational load (C,D) for the major (A,C) and minor (B,D) arrangements for different sized inversions. Color indicates presence (red) or absence (blue) of gene conversion. Each dot represents a single run.

**Figure S6.** Gene Conversion exponentially affects mutation accumulation and mutational load of the arrangements. (A) Boxplot showing the number of deleterious mutations accumulated in the major arrangement after 500,000 generations. Overlain points represent single runs where haplotype structuring did not occur (red) or did occur (blue). (B) Boxplot showing the number of deleterious mutations accumulated in the minor arrangement after 500,000 generations. Overlain points represent single runs where haplotype structuring did not occur (red) or did occur (blue). (C) Boxplot showing the mutational load of the major arrangement after 500,000 generations. Overlain points represent single runs where haplotype structuring did not occur (red) or did occur (blue). (D) Boxplot showing the log fitness the minor arrangement after 500,000 generations. Overlain points represent single runs where haplotype structuring did not occur (red) or did occur (blue). Five points, which were zero due to R’s internal cutoff, were replaced by 1 × 10^−7^.

**Figure S7.** Distribution of proportion of effectively neutral alleles among fixed mutations with (B,D) and without (A,C) gene conversion. Orange corresponds to mutations fixed in minor arrangement, cyan to mutations fixed in the major arrangement, pink to the average of mutations fixed in either the major or minor arrangement (i.e. alleles with an FST of 1), green to mutations that have fixed in the inverted region (i.e. fixed in both arrangements), and black to mutations that have fixed in the collinear region (chromosomes 2 and 3). The dashed black line indicate the proportion of new mutations that are effectively neutral, and the red dashed line corresponds to the proportion of effectively neutral mutations that fixed during the burn-in.

**Figure S8.** Density distribution of selective coefficient (log scale) of deleterious mutation with a Fst of 1 between the two arrangements. The red line indicates s=1/2N; to the left mutation are effectively neutral. A) All deleterious mutations within the inverted region, B) all deleterious mutations private to and fixed in the minority arrangement and C) all deleterious mutations private to and fixed in the majority arrangement.

**Figure S9.** Fitness distributions as a function of time reveal bimodality of the fitness of the homokaryotype. The different panels correspond to the fitness distribution of A) the whole population, B) the inversion homokaryotype, C) the heterokaryotype and D) the standard homokaryotype. The color indicates how many individuals share a given fitness values (on a log scale).

**Figure S10.** Fitness distributions as a function of time reveals that haplotype structuring happens in the absence of the initial heterozygote advantage (s_HET_=0) in the major arrangement. The different panels correspond to the fitness distribution of A) the whole population, B) the inversion homokaryotype, C) the heterokaryotype and D) the standard homokaryotype. The color indicates how many individuals share a given fitness value (on a log scale).

**Figure S11.** Formation of haplotype structuring in a model without an inversion. We consider a chromosome without an inversion but sharing the same properties than our inversion model (see methods for details) and determine the combination of crossing over and gene conversion rate where we observe haplotype structuring in at least 1 of 10 replicates (in red; black indicates that haplotype structuring was not observed). The × and Y axis corresponds to the relative values of crossing over and gene conversion rate compared to the main simulations.

**Figure S12.** Haplotype structuring when more than two haplotype clusters emerge in an arrangement. Panels A to D display the fitness distributions of A) the whole population, B) the homokaryotype for the inverted arrangement, C) the heterokaryotype and D) the homokaryotype for the standard arrangement. Panels A to D are similar to Figure S8 but for a different simulation run. Panel E) and F) corresponds to the allelic content of the inverted (E) and standard arrangement (F) at generation 500,000. Each horizontal line represents a haplotype in the population and each vertical line represents a genomic position. Yellow denotes that an individual possesses the derived allele and blue the ancestral one.

