## Supplemental Tables for "Deleterious Mutation Accumulation and the Long-Term Fate of Chromosomal Inversions"

### Supplemental Materials

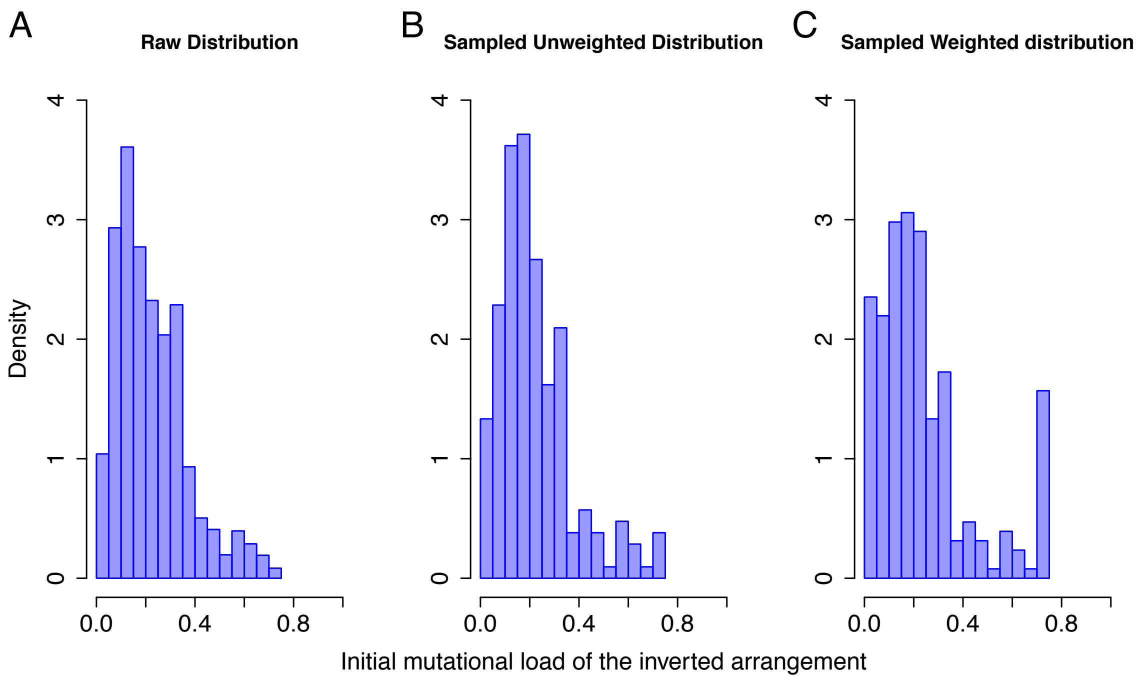


**Figure S1.**

Density distribution of the initial mutational load. A) the mutational load in the whole population at the end of the burn-in. B) the mutational load of the inverted arrangement in the haplotypes we selected (200 random plus the 4 best and the 4 worst and one close to the median). C) the mutational load of the inverted arrangement after correcting for the number of simulations done per haplotype. This figure illustrates that we do not always have the same number of simulations for each datapoint in Figure 1.

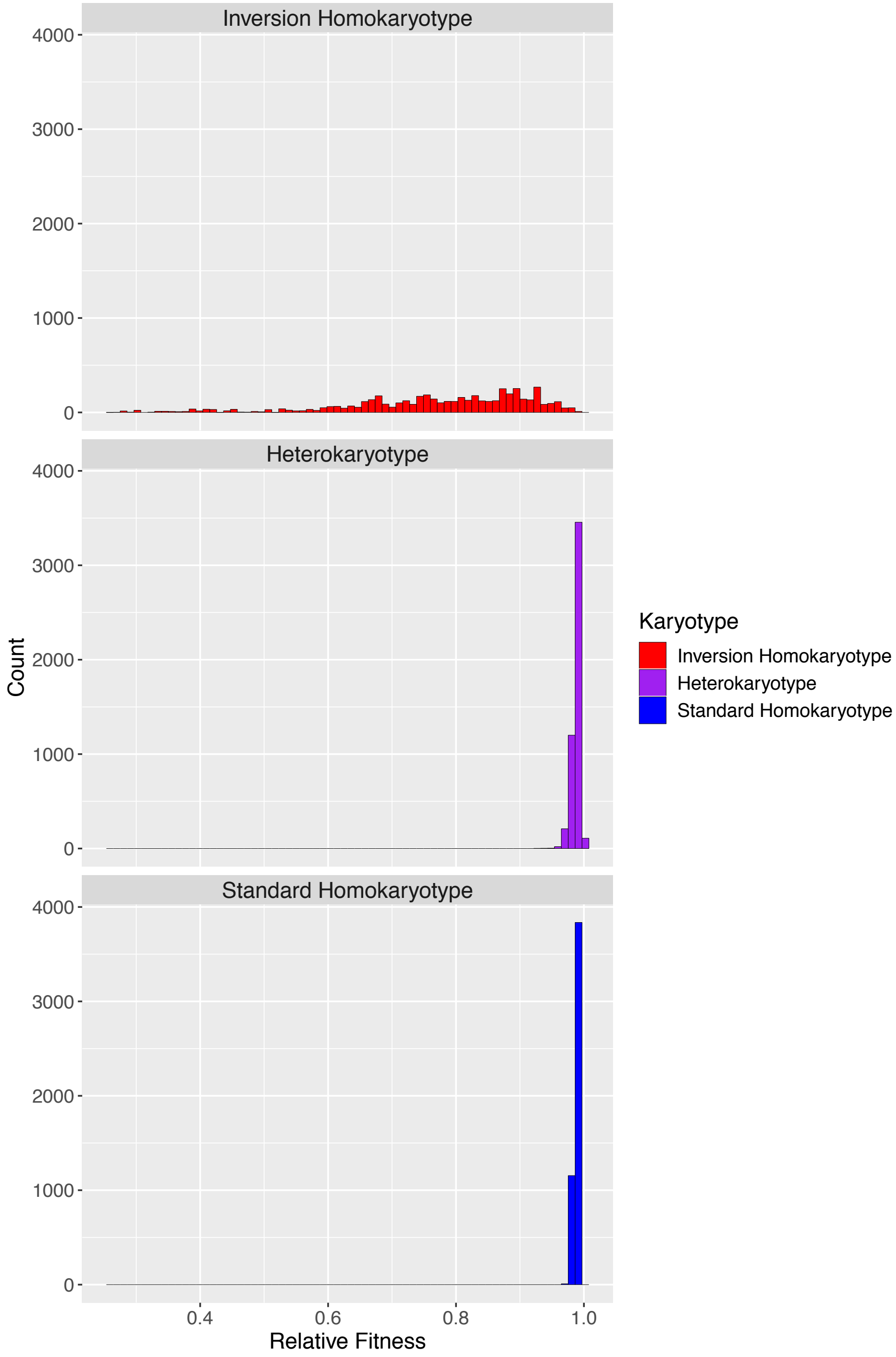


**Figure S2.**

Distribution of the initial relative fitnesses of all 3 karyotypes when an inversion occurs in any haplotype in a population.

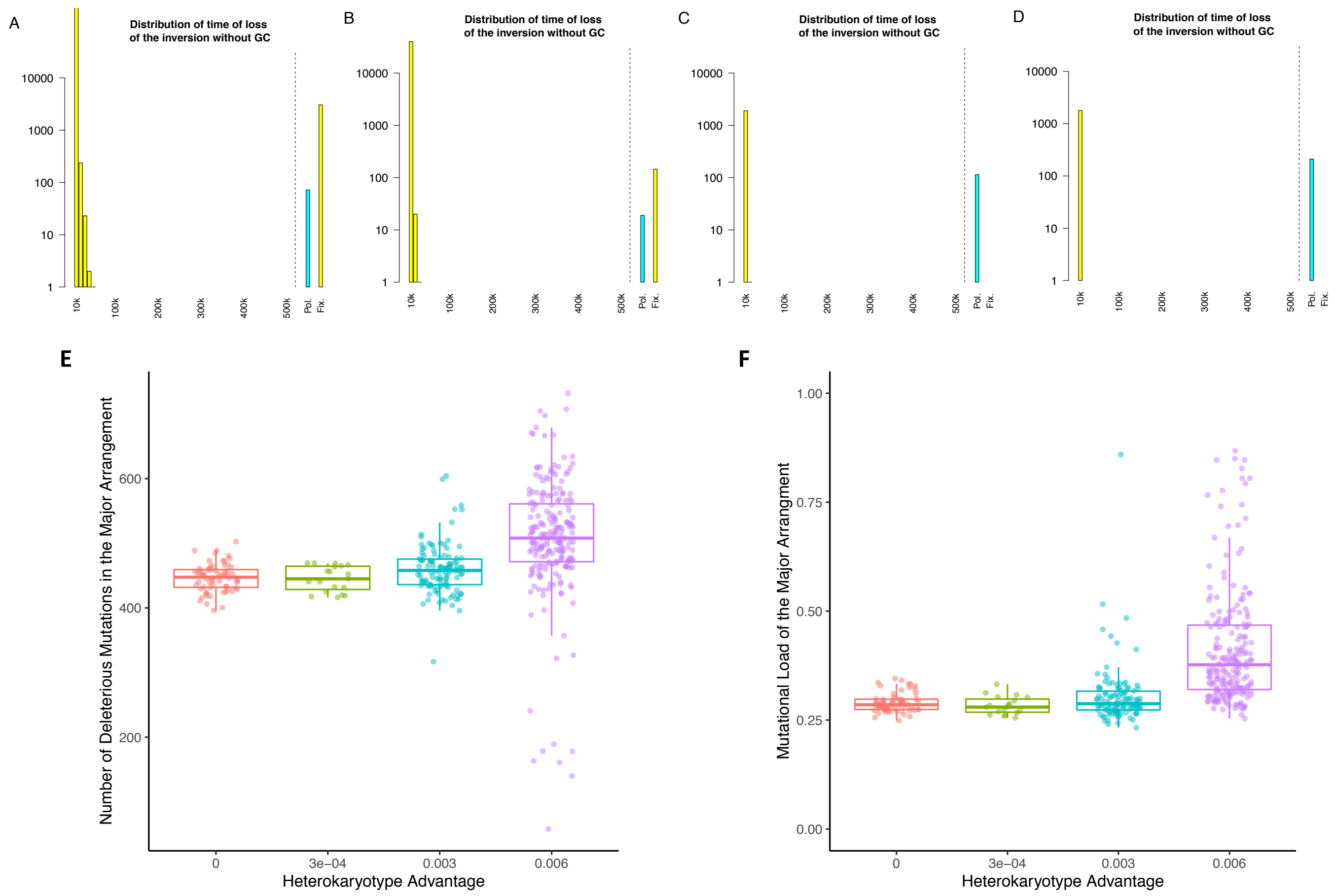


**Figure S3.**

Effects of the added heterokaryotype advantage. (A) Distribution of the time of loss of the inversion at $s_{HET}$=0. The number of simulations that remained polymorphic (cyan) or fixed (yellow) are indicated specifically to the right of the dashed line. (B) Distribution of the time of loss of the inversion at $s_{HET}$=0.0003. The number of simulations that remained polymorphic (cyan) or fixed (yellow) are indicated specifically to the right of the dashed line. (C) Distribution of the time of loss of the inversion at $s_{HET}$=0.003. The number of simulations that remained polymorphic (cyan) or fixed (yellow) are indicated specifically to the right of the dashed line. (D) Distribution of the time of loss of the inversion at $s_{HET}$=0.006. Simulations that remained polymorphic (cyan) or fixed (yellow) are indicated specifically to the right of the dashed line. (C) Mutation accumulation in the major arrangement under $s_{HET}$=0 (red), $s_{HET}$=0.0003 (green), $s_{HET}$=0.003 (cyan), and $s_{HET}$=0.006 (purple). Each dot represents a single run that ended at generation 500,000. (D) Mutation accumulation in the major arrangement under $s_{HET}$=0 (red), $s_{HET}$=0.0003 (green), $s_{HET}$=0.003 (cyan), and $s_{HET}$=0.006 (purple). Each dot represents a single run that ended at generation 500,000.

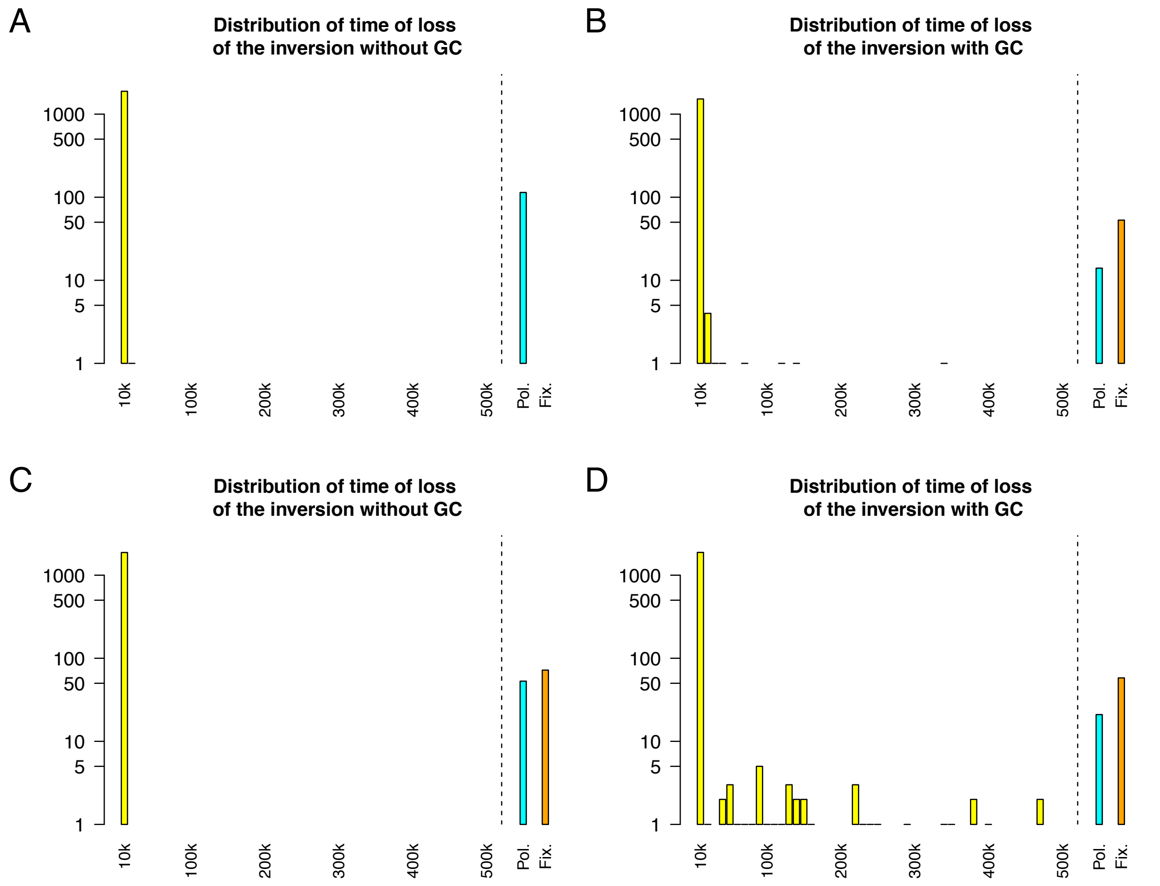


**Figure S4.**
Distribution of the time of loss of the inversion at different population sizes. For A and B N=25,000 and for C and D N=5,000. A and C show simulations run without gene conversion and C and D show simulations with gene conversion added. All plots show distribution of the time of loss of the inversion. Simulations that remained polymorphic (cyan) or fixed (yellow) are indicated specifically to the right of the dashed line.

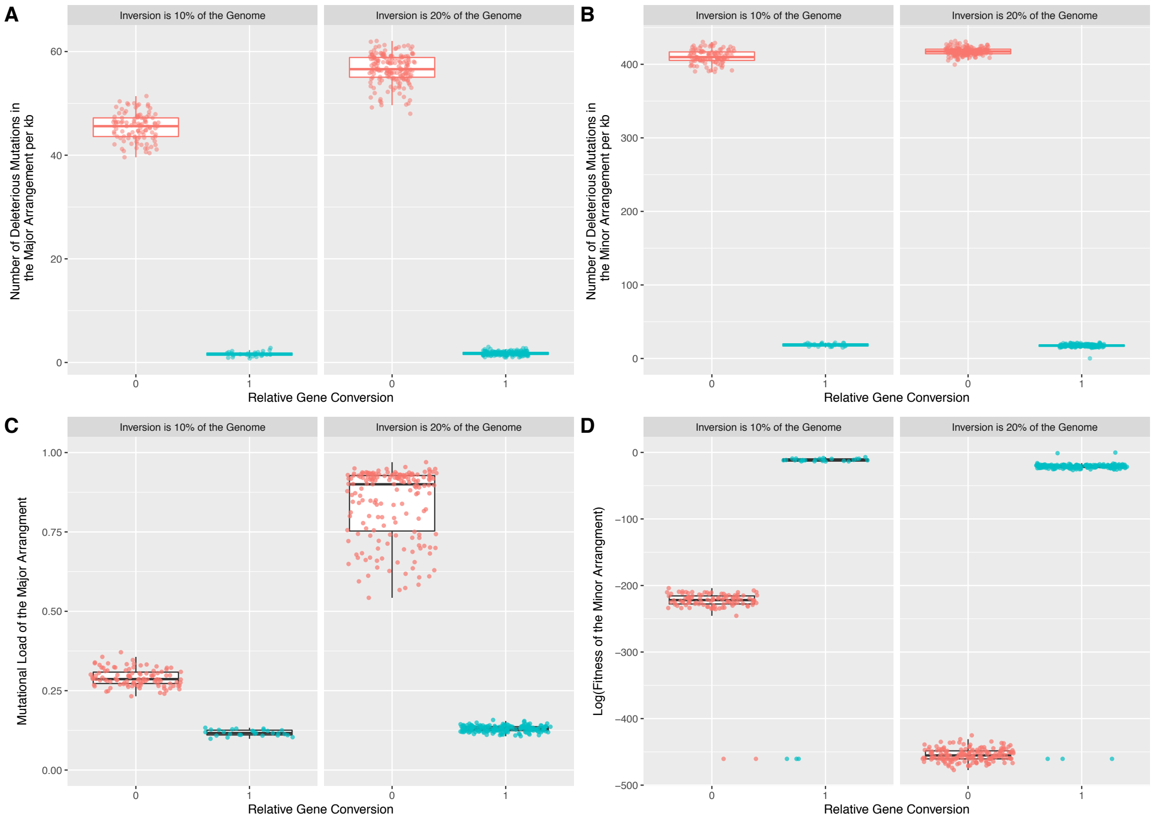


### Figure S5.

Mutation accumulation (A,B) and Mutational load (C,D) for the major (A,C) and minor (B,D) arrangements for different sized inversions. Color indicates presence (red) or absence (blue) of gene conversion. Each dot represents a single run.

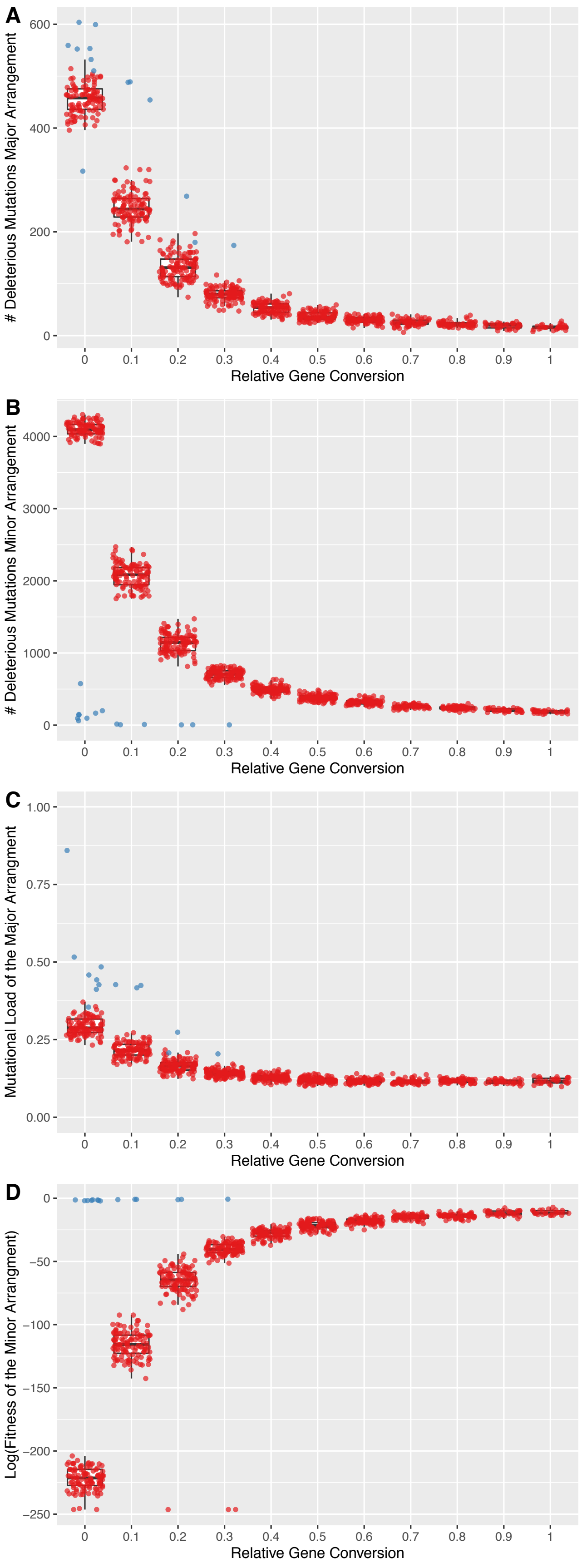
A
### Deleterious Mutations Major Arrangement
600
400
200
0
0
0.1
0.2
0.3
0.4
0.5
0.6
0.7
0.8
0.9
1
Relative Gene Conversion
B
### Deleterious Mutations Minor Arrangement
4000
3000
2000
1000
0
Relative Gene Conversion
C
Mutational Load of the Major Arrangment
1.00
0.75
0.50
0.25
0.00
Relative Gene Conversion
D
Log(Fitness of the Minor Arrangment)
0
−50
−100
−150
−200
−250
Relative Gene Conversion

**Figure S6.**

Gene Conversion exponentially affects mutation accumulation and mutational load of the arrangements. (A) Boxplot showing the number of deleterious mutations accumulated in the major arrangement after 500,000 generations. Overlain points represent single runs where haplotype structuring did not occur (red) or did occur (blue). (B) Boxplot showing the number of deleterious mutations accumulated in the minor arrangement after 500,000 generations. Overlain points represent single runs where haplotype structuring did not occur (red) or did occur (blue). (C) Boxplot showing the mutational load of the major arrangement after 500,000 generations. Overlain points represent single runs where haplotype structuring did not occur (red) or did occur (blue). (D) Boxplot showing the log fitness the minor arrangement after 500,000 generations. Overlain points represent single runs where haplotype structuring did not occur (red) or did occur (blue). Five points, which were zero due to R's internal cutoff, were replaced by $1 \times 10^{-7}$.

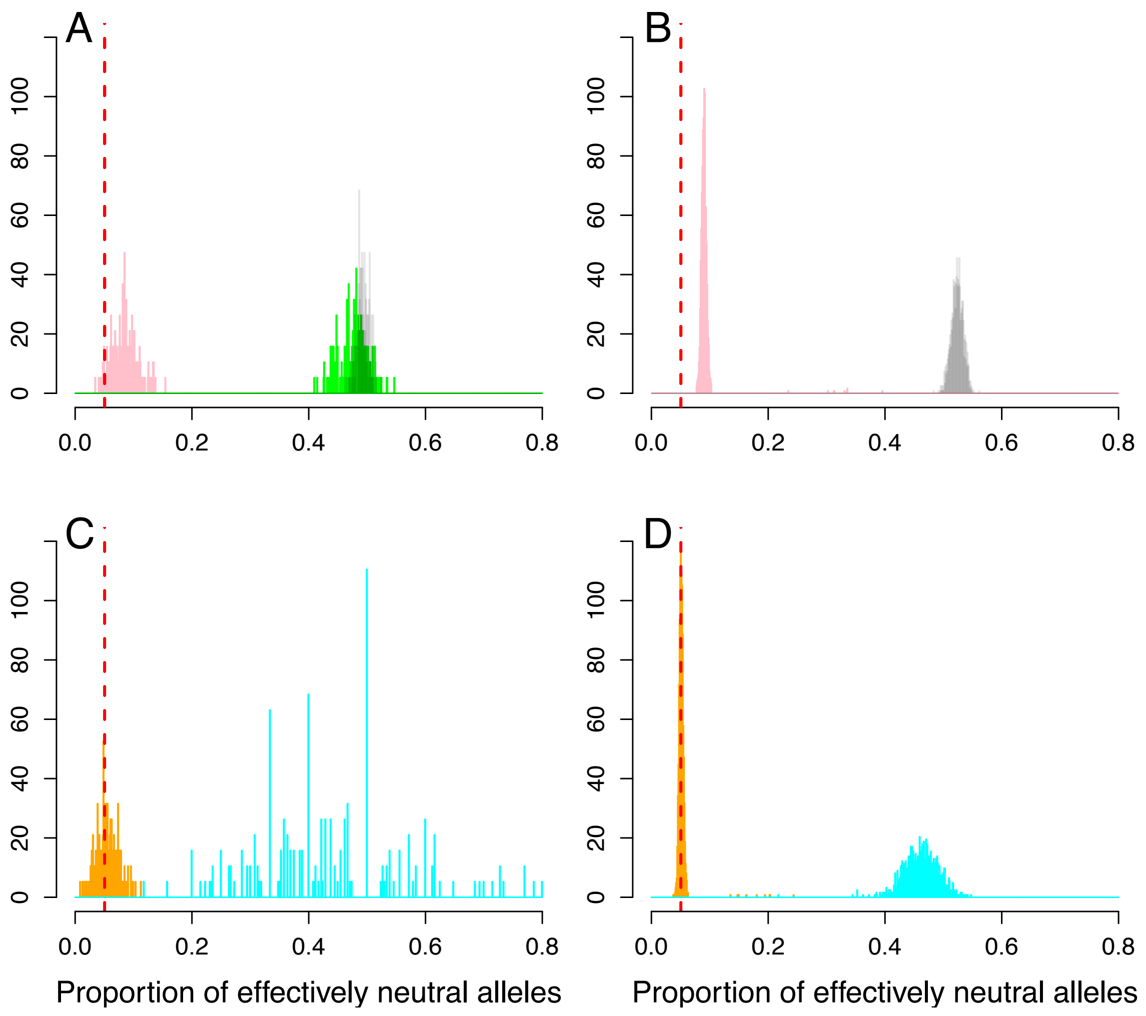


**Figure S7.**

Distribution of proportion of effectively neutral alleles among fixed mutations with (B,D) and without (A,C) gene conversion. Orange corresponds to mutations fixed in minor arrangement, cyan to mutations fixed in the major arrangement, pink to the average of mutations fixed in either the major or minor arrangement (i.e. alleles with an FST of 1), green to mutations that have fixed in the inverted region (i.e. fixed in both arrangements), and black to mutations that have fixed in the collinear region (chromosomes 2 and 3). The dashed black line indicate the proportion of new mutations that are effectively neutral, and the red dashed line corresponds to the proportion of effectively neutral mutations that fixed during the burn-in.

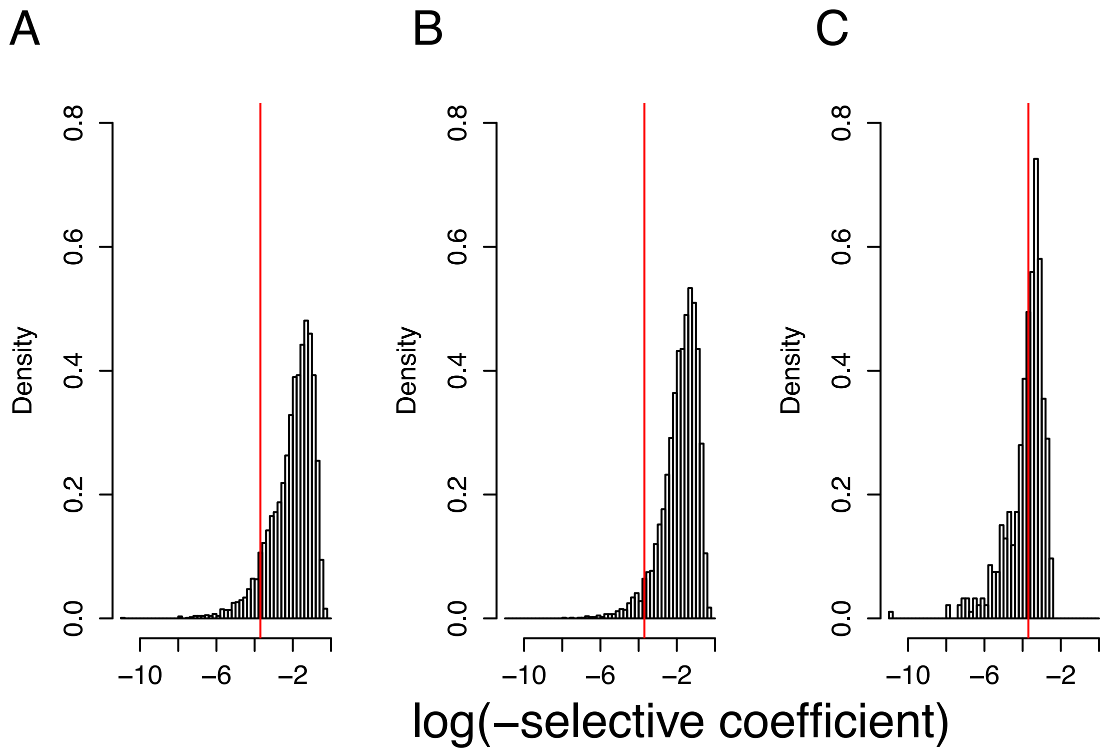


**Figure S8.**
Density distribution of selective coefficient (log scale) of deleterious mutation with a Fst of 1 between the two arrangements. The red line indicates s=1/2N; to the left mutation are effectively neutral. A) All deleterious mutations within the inverted region, B) all deleterious mutations private to and fixed in the minority arrangement and C) all deleterious mutations private to and fixed in the majority arrangement.

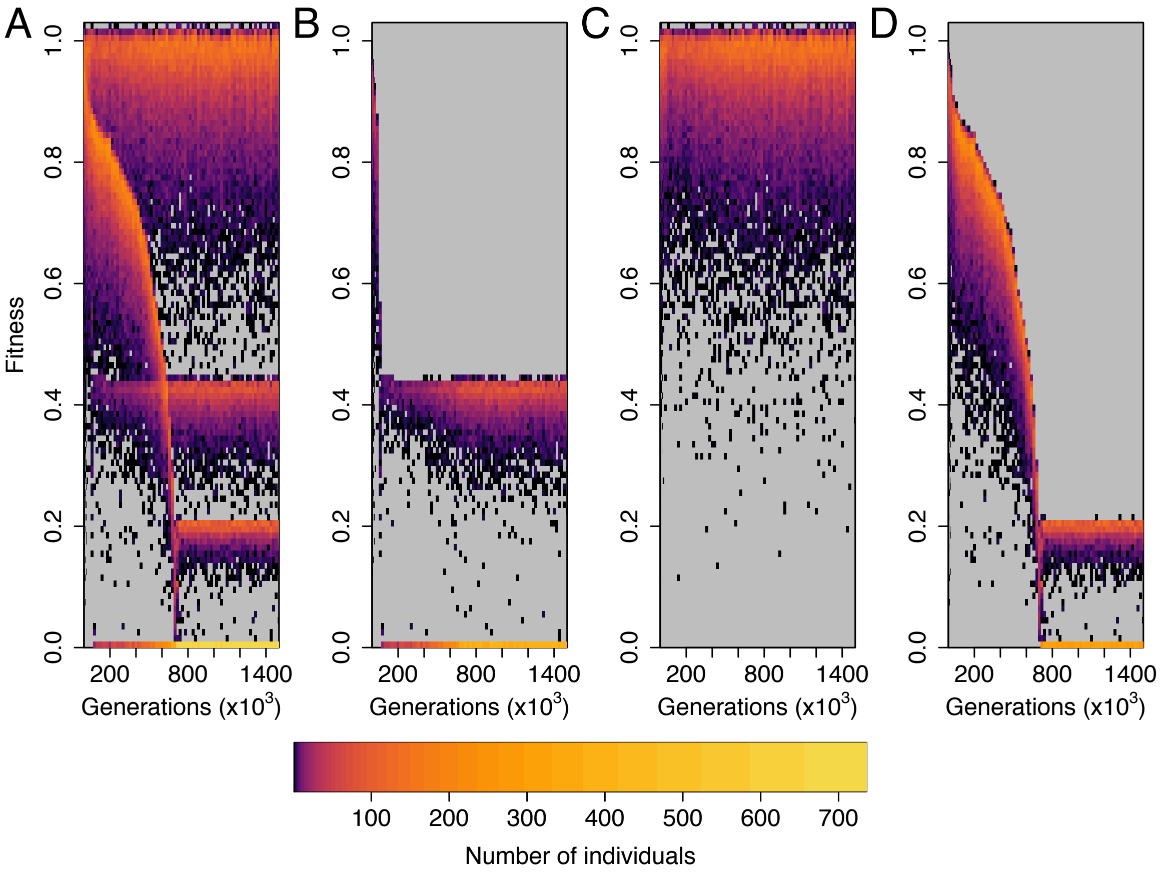


**Figure S9.**

Fitness distributions as a function of time reveal bimodality of the fitness of the homokaryotype. The different panels correspond to the fitness distribution of A) the whole population, B) the inversion homokaryotype, C) the heterokaryotype and D) the standard homokaryotype. The color indicates how many individuals share a given fitness values (on a log scale).

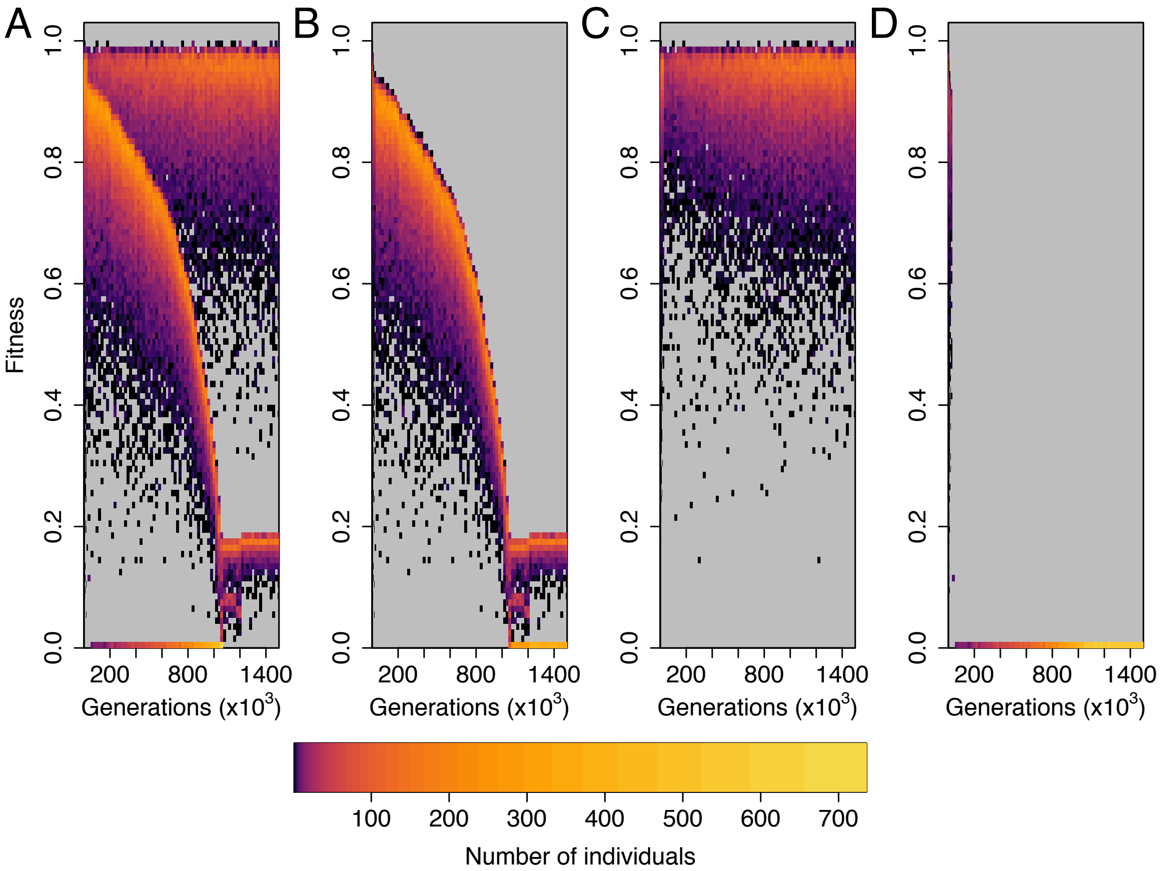


**Figure S10.**
Fitness distributions as a function of time reveals that haplotype structuring happens in the absence of the initial heterozygote advantage ($s_{HET}$=0) in the major arrangement. The different panels correspond to the fitness distribution of A) the whole population, B) the inversion homokaryotype, C) the heterokaryotype and D) the standard homokaryotype. The color indicates how many individuals share a given fitness value (on a log scale).

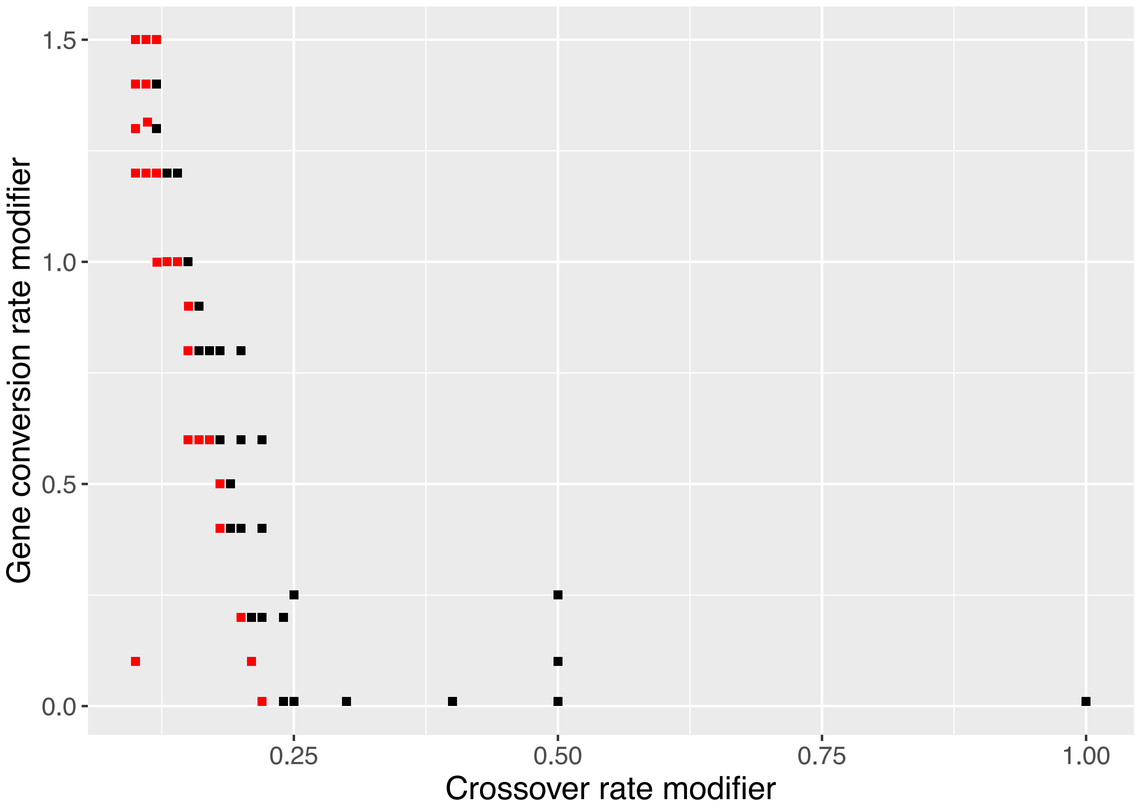


**Figure S11.**

Formation of haplotype structuring in a model without an inversion. We consider a chromosome without an inversion but sharing the same properties than our inversion model (see methods for details) and determine the combination of crossing over and gene conversion rate where we observe haplotype structuring in at least 1 of 10 replicates (in red; black indicates that haplotype structuring was not observed). The X and Y axis corresponds to the relative values of crossing over and gene conversion rate compared to the main simulations.

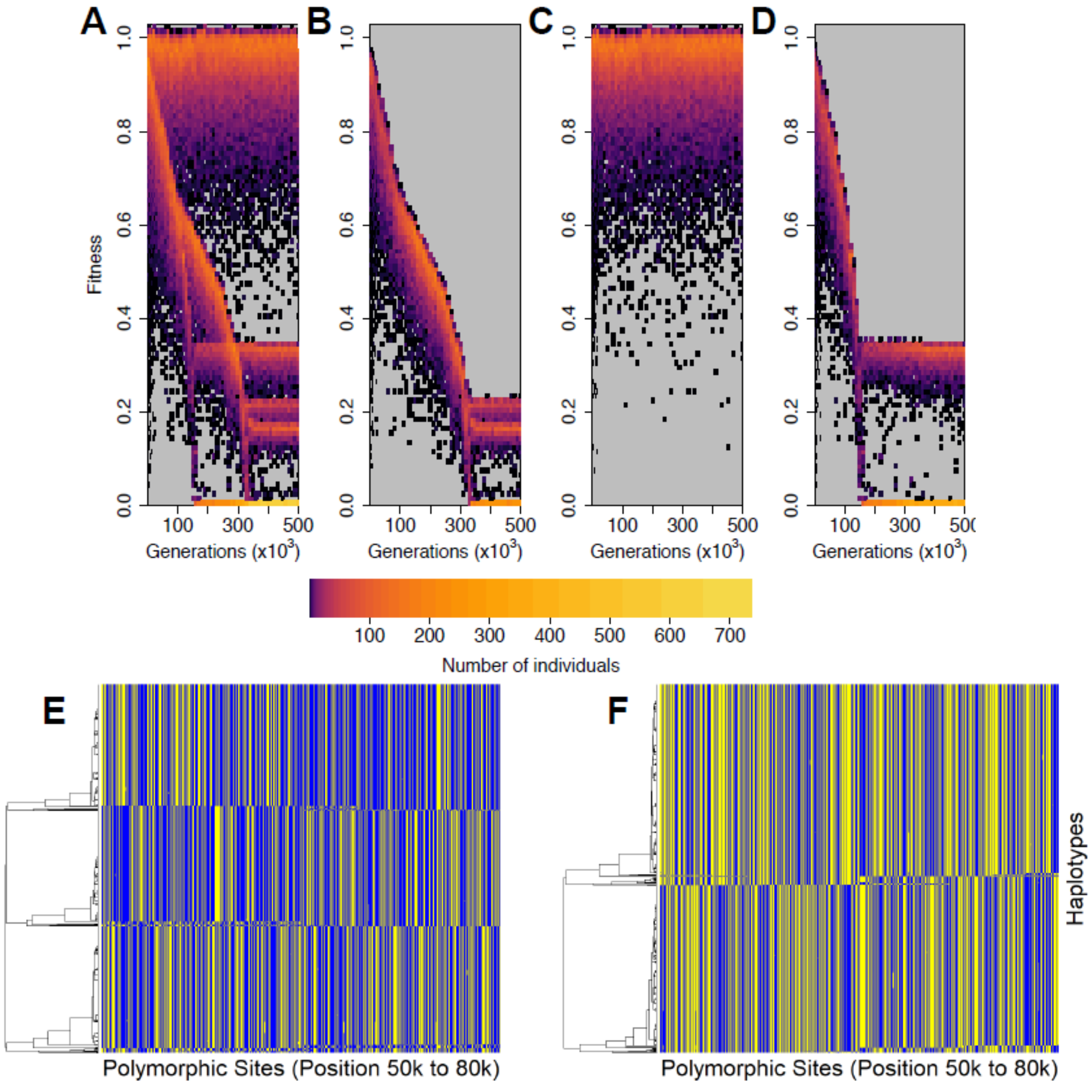


#### Figure S12.

Haplotype structuring when more than two haplotype clusters emerge in an arrangement. Panels A to D display the fitness distributions of A) the whole population, B) the homokaryotype for the inverted arrangement, C) the heterokaryotype and D) the homokaryotype for the standard arrangement. Panels A to D are similar to Figure S8 but for a different simulation run. Panel E) and F) corresponds to the allelic content of the inverted (E) and standard arrangement (F) at generation 500,000. Each horizontal line represents a haplotype in the population and each vertical line represents a genomic position. Yellow denotes that an individual possesses the derived allele and blue the ancestral one.
