## Supplemental Text for "Deleterious Mutation Accumulation and the Long-Term Fate of Chromosomal Inversions"

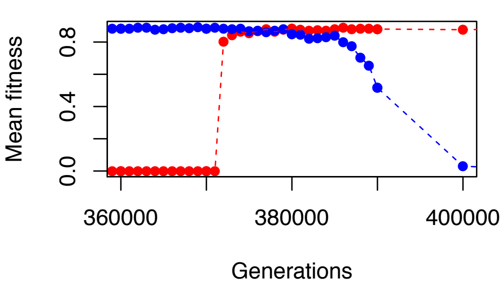


(a) Mean fitness of the homokaryotypes

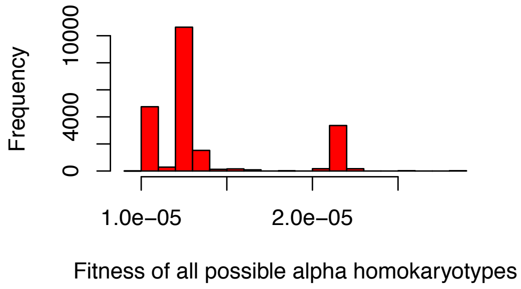


(b) Distribution of homokaryotype fitness at generation 370000 for the inverted arrangement

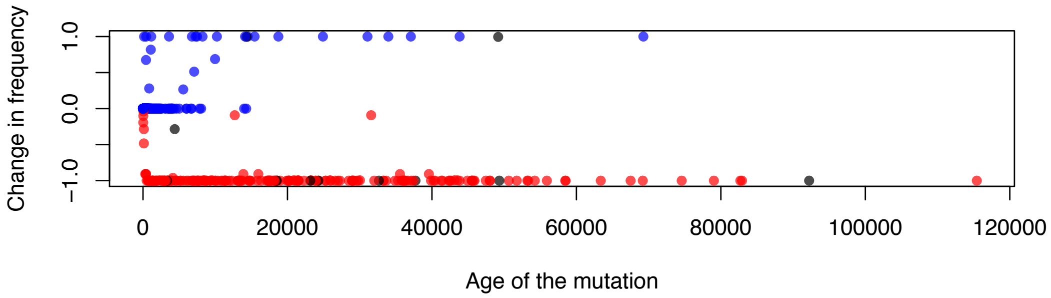


**Figure 1.** Fitness recovery of a previous inviable homokaryotype. (a) Mean fitness for the inverted (red) and standard (blue) homokaryotype. (b) Distribution of the fitness of all possible inverted homokaryotypes at generation 370 000 (before the increase of the inverted homokaryotype mean fitness). (c) Change in frequency of all mutations in the inverted region that are present at generation 370000. The X axis indicates the age of the mutations in generation 370000 and the color indicates if the mutation at generation 370000 was private to the inverted arrangement (red), private to the standard arrangement (blue) or shared between the two arrangements (black).

Once the fitness of an homokaryotype is close to zero, it is unable to reproduce. The homokaryotype almost never recovered from this state. Among all simulations, a permanent recovery was observed only once in a run with one of the four fittest haplotypes at 90% of our normal gene conversion rate. In this case, illustrated in Figure 1, the fitness of the minor homokaryotype recovered to the same level as the major homokaryotype. After a period of similar fitness, the fitness of the other homokaryotype decreased (in that case the standard arrangement) due to mutation accumulation, as illustrated in Figure 1, panel A.

There are two possible explanations for the recovery of the fitness of the minor haplotype: (1) there was a rare haplotype containing few mutations which swept through the arrangement subpopulation or (2) a minor haplotype was relieved from its mutations by a rapid successful introgression from the major arrangement into the minor arrangement. Either scenario is unlikely which explains why we only observed this once. The first scenario requires the existence of a mostly mutation-free haplotype that is maintained at low frequency for a long time; this is unlikely given the number of copies of the minor arrangement. The second scenario requires the replacement of the allelic content of the minor arrangement with the allelic content of the major arrangement in a relatively short time to prevent further degradation due to mutation accumulation. Introgression from one arrangement to the other can only happen in our model due to gene conversion events that have an average length of 500 bp. Replacement of the minor allelic content by the major allelic content therefore requires many gene conversion events happening in the same haplotype in a relatively short period of time.

To distinguish between the two scenarios described above, we first examined the distribution of the fitness of all possible minor arrangement homokaryotypes in the last sampled generation before the fitness increase (Figure 1, panel B). All possible generated homokaryotypes had fitnesses approaching zero, meaning that we can safely reject our first hypothesis. To confirm the second hypothesis, we looked at different mutations that were present in the population before and after the fitness increase of the minor arrangement. We illustrate the change in frequency of all mutations in the inverted region that existed at generation 370 000 (Figure 1 panel C). The transitional period is characterized by the loss of almost all mutations private to the minor arrangement (in red) while many mutations previously private to the major arrangement (in blue) fixed (or almost fixed) in the minor arrangement. This indicates that the allelic content of the minor arrangement was replaced by introgression from the major arrangement.
